# Planar, organoid-independent iPSC-derived small intestinal epithelium from celiac disease patient for studying cytokine-induced changes

**DOI:** 10.64898/2026.03.12.710771

**Authors:** Pauliina Kukkoaho, Maria Annala, Marika Kosunen, Helka Kaunisto, Karoliina Tanner, Fatima Siddique, Ella Soltin, Niyati Kandikanti, Susanna Kaksonen, Katarzyna Leskinen, Päivi Saavalainen, Juha Kesseli, Matti Nykter, Katriina Aalto-Setälä, Katri Kaukinen, Katri Lindfors, Kati Juuti-Uusitalo

**Affiliations:** Faculty of Medicine and Health Technology, Tampere University, 33014 Tampere; Folkhälsan, 00250 Helsinki; University of Helsinki, 00290 Helsinki; Heart Hospital, Tampere University Hospital, 33521 Tampere, Finland; Department of Internal Medicine, Tampere University Hospital, 33521 Tampere, Finland

**Keywords:** cytokine, interferon gamma, tumor necrosis factor alpha, induced pluripotent stem cell, iPSC, cytokine, celiac disease, small intestinal epithelial cell

## Abstract

**Background & Aims:** Celiac disease is a wheat-induced immune-mediated enteropathy characterized by cytokine, especially IFNγ upregulation. Intestinal organoid models for adult stem cell–based celiac disease exist, but planar iPSC based small intestinal epithelial models derived from celiac disease patients that would allow direct assessment from both sides of the epithelium have been lacking. We aimed to bridge this gap by setting up a two-dimensional *in vitro* model based on small intestinal epithelial cells derived from iPSC from celiac disease patient.

**Methods:** IPSCs from celiac disease patient and control patient were directly differentiated into small intestinal epithelium on Transwell inserts without intermediate organoid phase or replating. Model maturity and celiac disease specific cytokine responses were analysed using immunofluorescence, transcriptomic profiling, and LDH cytotoxicity assay.

**Results:** Two-dimensionally cultured **c**eliac disease derived line and control patient derived iPSCs matured similarly towards intestinal epithelium which expressed enterocyte specific PEPT1 and EpCAM. To the cytokine stimuli, celiac and healthy iPSC-SIECs responded by increasing the expression IRF1, ICAM1 and IDO1, and several other known cytokine responsive genes.

**Conclusions:** These findings suggest that the 2D iPSC celiac disease small intestinal epithelium model represent an appropriate platform for studying inflammation-associated enteropathies, such as celiac disease.

## 1. Introduction

Celiac disease (CeD) is a common immune-mediated enteropathy, which affects approximately 1%–2% of the population (1,2). The onset of CeD requires both gluten ingestion and a genetic predisposition (1,2). Out of all CeD patients, 95% are carriers of human leukocyte antigen (HLA) risk haplotypes HLA-DQ2 or HLA-DQ8 (3–5). In 2% of the carriers of CeD-predisposing HLA-DQ haplotypes (1,2), incompletely digested gluten, after tissue transglutaminase deamidation, is able to elicit the activation of an immune response (5). The adaptive immune reaction leading to the onset of CeD has been characterized in detail, but the direct effects of gluten and the effects of the microbiome on the epithelium and CeD onset remain insufficiently understood (6–8).

Research on the CeD-affected intestinal epithelium is still often carried out using immortalized cell lines, such as Caco-2, which have a cancerous origin (9,10). Biopsy-derived intestinal adult stem cells (ACS) from CeD patients that are grown within an extracellular matrix to form organoids resemble their native counterparts in that they carry CeD genetics and have the morphology and cellular functionality of intestinal epithelial cells (10–18). These three-dimensionally cultured organoids, however, have a closed, apically inaccessible structure that is not optimal for studying the effects of environmental stimuli, such as nutrient uptake—e.g., that of gliadin—or microbial stimuli (19,20). Three dimensional ACS derived organoids must be opened and transitioned to two dimensional (2D) planar cultures for use in transport studies (10–12,14,15). Furthermore, when aiming to develop autologous multi cellular epithelial models, it is important to note that ASCs inherently include the same cell types present in an intestinal biopsy and that their intestinal stem cells can only differentiate into intestinal epithelial lineages. In contrast, induced pluripotent stem cells (iPSCs) are capable of differentiating into a wide range of cell types, enabling the creation of more complex autologous multicell models (19).

IPSCs can be differentiated within an extracellular matrix to form organoids (21), and they can be similarly opened to form planar cultures (22). However, there is no need to differentiate iPSCs into small intestinal epithelial cells (SIECs) via organoids, but the differentiation can be carried out directly on a planar surface in 2D, in an apically accessible format (23–25). In this study, we chose to bypass the conventional 3D organoid culture step and instead directly differentiate iPSCs into planar SIECs using a protocol previously optimized in our laboratory (25), based on methods developed by the Matsunaga and Mizuguchi groups (23,24,26,27).

When developing preclinical models for certain disease entities, it is vitally important to test how cells respond to disease-specific stimuli. In CeD the intestinal mucosa expression of several cytokines is elevated (28) and further increased after a gluten challenge (29). The interferon γ (IFNγ) expression has been shown to be increased more than 1,000-fold (30). Tumor necrosis factor α (TNFα) expression is upregulated in the CeD-affected intestine (29) but also in another enteropathy, namely inflammatory bowel disease (31,32).

To the best of our knowledge, there have been no earlier publications involving a model utilizing small intestinal epithelial cells (SIECs) derived from iPSCs of CeD patient (i.e., iPSC-SIEC model). The aim of the present study was to establish an iPSC-SIEC model that bypasses organoid differentiation stage, characterize the differentiation status, and as a proof-of-concept evaluate transcriptomic effects of CeD-specific cytokine stimuli.

## 2. Materials and Methods

### 2.1. iPSC culture

Here four iPSC lines derived by means of viral transduction in iPSC core (Tampere University, Tampere, Finland) (33,34) were used. The genotyping of iPSC cell lines was carried out using Olerup SSP® DQ low resolution kit (Olerup SSP AB, Stockholm, Sweden) with protocol previously published in Kurppa et al. (35). The UTA.10606.EURCAp from celiac patient, UTA.08102.EURCCS from type 1 diabetic patient carried of high-risk celiac-predisposing gene HLA-DQ2, whereas the 04602.WT iPSC line (25) from non-celiac patient and the UTA.10802.EURCCS line (33) from type 2 diabetic patient were carriers of HLA-Q7 and HLA-DQ4, respectively. At the time of the experiment, the karyotypes of the iPSC lines were tested to be normal (Fimlab Laboratories, Tampere, Finland). The iPSCs were maintained on Geltrex-coated (Gibco; Thermo Fisher Scientific, Waltham, MA, USA; diluted 1:100) Cell-Bind-treated 6-well plates (Corning, Corning, NY, USA) in mTeSR1 medium (StemCell Technologies, Vancouver, Canada) supplemented with penicillin-streptomycin (P/S; Gibco). In maintaining the culture, the cells were passaged using Versene (Gibco) when approximately 90% confluent and seeded at 3-4*10^4^ cells/cm^2^. In Table 1. the times when the cells were accessed, from the thawing to the end of the cytokine experiment.

**Table 1.** Times when cells were accessed. Time starts with the thawing of the iPSCs and finishes to the end of the cytokine stimuli experiment.

| CeD | HC | T1D | T2D |
|---|---|---|---|
| 30/06/2022 - 11/08/2022 | 16/06/2022-28/07/2022 | 06/07/2022-11/08/2022 | 06/07/2022-11/08/2022 |
| 08/07/2022-17/08/2022 | 06/07/2022-11/08/2022 | 06/07/2022-17/08/2022 | 06/07/2022-17/08/2022 |
| 06/06/2024 - 11/07/2024 | 25/08/2023 - 26/09/2023 | 14/07/2022 - 24/08/2022 | 14/07/2022-24/08/2022 |
| 14/06/2024 – 17/07/2024 | 19/09/2023 - 25/10/2023 |  | 25/08/2023 - 26/09/2023 |
| 26/06/2024-01/08/2024 | 06/06/2024 - 11/07/2024 |  | 19/09/2023 – 25/10/2023 |
| 02/07/2024 07/08/2024- | 14/06/2024 – 17/07/2024 |  |  |
| 12/05/2026 - 17/06/2026 | 26/06/2024-01/08/2024 |  |  |
| 27/08/2026 – 29/09/2026 | 02/07/2024 - 07/08/2024 |  |  |
|  | 13/08/2026-17/09/2026 |  |  |

### 2.2. In vitro differentiation into small intestinal epithelial cells

The differentiation towards small intestinal epithelial cells was done as described previously in Saari et al 2022 (25). Briefly, iPSCs were seeded (1-1.15x10^5^ cells/cm^2^) onto human recombinant laminin-coated PET cell culture inserts (LN111 1.8 μg/cm^2^, LN511 0.6 μg/cm^2^ Biolamina, Stockholm, Sweden; 1 μm pores Sarstedt, Nümbrecht, Germany) and cultured in mTeSR1 for 24 h. Definitive endoderm (DE) differentiation was initiated by switching to RPMI1640 supplemented with GlutaMax (both Gibco, Thermo Fisher Scientific), Activin A (100 ng/ml, Miltenyi Biotec, Bergisch Gladbach, Germany), and P/S for 72 h. During the first 24 h, CHIR99021 (2.5μM, Tocris Biosciences, Bristol, UK) and B27-vitamin-A (Gibco) were added, followed by supplementation with 0.2% and 2% KnockOut Serum Replacement (KO-SR, Gibco) during the second and third 24-h periods, respectively. After DE, the cultures were gently washed once with 1X DPBS.

Posterior definitive endoderm (PDE) differentiation was carried out for 96 h as described previously in Saari et al (25), following a DPBS wash. Cells were cultured in high-glucose DMEM, NEAA (both (Thermo Fisher Scientific), LY2090314 (20 nM, MedChem Express, Monmouth Junction, NJ, USA), GlutaMax, P/S, and 10% KO SR, with medium changes every two days. After PDE cultures were gently washed with DPBS. Maturation to SIEC was subsequently performed over 21 days in DMEM/F-12 containing 20% KO-SR, NEAA, GlutaMax, B27 (-vitamin-A), N2, and P/S (both Gibco), supplemented with rhWnt3A, rhEGF, rhIGF (all from R&D Systems), dexamethasone (Tocris Biosciences), rhR-spondin-3, rhNoggin (both Miltenyi Biotech), and SB202190 Merck, Sigma-Aldrich). Medium was refreshed every three days, completing a total differentiation period of 28 days. Only cultures retaining medium in the apical chamber were used for stimulation experiments, which were initiated on day 26. During the cell culture, cells were imaged with phase contrast microscope using Zeiss Axiovert A1 5x LD A-Plan lens and Zeiss Axiocam ERc 5s camera (Carl Zeiss Microscopy GmbH, Göttingen, Germany).

### 2.3. Cytokine stimulation

Cytokine stimulation protocols were adapted from previous studies. IFNγ stimulation was modified from Workman et al. (2020) (36), Van De Walle et al. (2010) (37), and Zolotarevsky et al. (2002) (38), with concentrations of 10 ng/ml and 50 ng/ml selected based on Workman et al. (2020) (36) and Van De Walle et al. (2010) (37), respectively. The 48-h stimulation Foster et al. (2009) (39), basolateral stimulation as described by Van De Walle et al. (2010) (37). TNFα stimulation concentration (10 ng/ml) on Al-Sadi et al. (2013) (40), with stimulation duration and basolateral application according to Ma et al. (2005) [40] and Zolotarevsky et al. (2002) (38).

Both cytokines (PeproTech, Thermo Fisher Scientific) were reconstituted according to the manufacturer’s instructions and diluted immediately prior to use. Stimulation was initiated on day 26 by adding induction medium to the basolateral chamber and was maintained for 48 h, with medium refreshed after 24 h. Samples (150 µl) were collected from both apical and basolateral compartments at 0, 24, and 48 h. Collected media were centrifuged (200 g, 5 min), and supernatants were stored at -80°C to preserve lactate dehydrogenase activity (41).

### 2.4. Transepithelial electrical resistance

Transepithelial electrical resistance (TEER) was measured with Millicell® ERS 3.0 Digital Voltohmmeter (Millipore/ Merck KGaA), Darmstadt, Germany) on day 25 (healthy control-patient-derived line) or day 26 CeD-derived line. TEER values were correlated with culture morphology assessed with Evident FLUOVIEWTM FV4000 laser scanning confocal imaging system from phalloidin and DAPI stained samples.

### 2.5. Immunofluorescence staining and imaging

Immunofluorescence staining was performed largely as described previously (Saari et al., 2022 (25)). Briefly, following stimulation, cells were washed with DPBS, fixed in 4% paraformaldehyde (Thermo Fisher Scientific, Waltham, MA, USA) for 10 min, permeabilized with 0.1% Triton X-100 for 10 min, and blocked with 3% bovine serum albumin for 1 h. Primary antibodies (Table 2) were diluted in 0.5% BSA and incubated for 1 h, followed by secondary antibodies (Table 2) supplemented with Alexa Fluor 633 Phalloidin (4 U/ml, Invitrogen, Thermo Fisher Scientific). Samples were washed between steps and mounted with Vectashield containing DAPI (Vector Laboratories, Burlingame, CA, USA).

**Table 2.** Antibodies used for immunofluorescence staining.

| **Primary Antibodies** |  |  |  |  |  |
| **Target** | **Target marker for** | **Manufacturer** | **Catalog** | **Host** | **Dilution** |
|---|---|---|---|---|---|
| Oct4 | pluripotency | R&D | AF1759 | goat | 1:300 |
| Nanog | pluripotency | R&D | AF1997 | goat | 1:300 |
| Sox17 | endoderm | R&D | AF1924 | goat | 1:300 |
| Foxa2 | endoderm | EMD Millipore | 07-633 | rabbit | 1:300 |
| Pept1 | enterocytes | Novus Biologicals | NBPI-92005 | rabbit | 1:100 |
| Villin | epithelial cells | Abcam | ab130751 | rabbit | 1:200 |
| E-cadherin | epithelial junctions | Abcam | ab231303 | mouse | 1:200 |
| EpCam | epithelial cells | CST | 2929S | mouse | 1:300 |
| **Secondary Antibodies** |  |  |  |  |  |
| **Target** | **Manufacturer** | **Wavelength** | **Cat** | **Host** | **dilution** |
| Goat IgG | Invitrogen | 568 | A11057 | Donkey | 1:500 |
| Rabbit IgG | Invitrogen | 488 | A21206 | Donkey | 1:500 |
| Mouse IgG | Invitrogen | 488 | A21202 | Donkey | 1:500 |
| Rabbit IgG | Thermo Scientific | 568 | A10042 | Donkey | 1:500 |

IPSC, DE, and PDE samples were stained and imaged directly in their culture vessels using an Olympus IX-51 epifluorescence microscope (Olympus Corporation, Tokyo, Japan) equipped with a Hamamatsu Orca Flash4.0LT+ sCMOS camera (Hamamatsu Photonics Europe GmbH, Herrsching am Ammersee, Germany). Images were colorized using Adobe Photoshop (Adobe Inc., San José, CA, USA).

SIEC cultures were grown on inserts; to reduce reagent use, membranes were detached from the insert holders and stained on glass slides using aqueous droplets.

Confocal imaging of anti-PEPT1 labeled samples was performed using an LSM800 laser scanning confocal microscope (Zeiss Carl AG, Oberkochen, Germany) with a Plan-Apochromat 63x/1.4 NA oil immersion objective. DAPI, Alexa Fluor 488, and Alexa Fluor 647 were excited at 405, 488, and 633 nm, respectively, and emission was detected using GaAsP-PMT detectors. Alexa Fluor 488 and 647 were acquired simultaneously, followed by sequential acquisition of the DAPI channel. Images were acquired without averaging using a 0.52 μs pixel dwell time, a 1 Airy unit pinhole, a 99 nm xy pixel size, and 200 nm z-steps over manually defined z-ranges.

Confocal imaging of Villin and/ or E-cadherin and/ or EpCAM labeled samples was performed using Olympus IX83 inverted microscope with Evident FLUOVIEWTM FV4000 laser scanning confocal imaging system (Evident Scientific, Tokyo, Japan) with an Olympus U Plan-Apochromat 60x/1.30 NA silicone immersion objective. DAPI, Alexa Fluor 488, Alexa Fluor 568 and Alexa Fluor 640 were excited at 405, 488, 561 nm and 640 nm, respectively. Emissions of 405 nm and 488 nm were detected with Broadband SilVIR, 561 nm and 640 nm with Red-shifted SilVIR detector. All channels were imaged separately. Images were acquired without averaging using a 0.52μs pixel dwell time, a 1 Airy unit pinhole, a 99 nm xy pixel size, and 300 nm z-steps over manually defined z-ranges.

Image stacks were deconvolved using Huygens Essentials (v25.04, Scientific Volume Imaging, The Netherlands) with the CMLE algorithm and a theoretical point spread function, using automatic background correction. Maximum-intensity projections (MIPs) of apical (XY) and orthogonal (XZ/YZ) planes were generated using a custom ImageJ2 (v2.16.0/1.54p) macro. Apical MIPs were generated over a 6μm region starting from the detected apical signal onset, while orthogonal MIPs were generated over a 1μm range centered on user-defined positions. Brightness and contrast were adjusted in ImageJ2 for optimal visualization, and all projections were manually inspected and corrected when necessary.

### 2.6. Gene expression analysis

#### 2.5.1. mRNA extraction

After each differentiation step and after cytokine stimuli, RNA was extracted from the cells similarly to the protocol in Saari et al 2022 (25) using the RNeasy Plus Mini Kit (Qiagen, Hilden, Germany) according to the manufacturer’s protocol. RNA concentration and purity were measured using a NanoDrop One spectrophotometer (NanoDrop Technologies).

#### 2.6.2. qRT-PCR

Reverse transcription was performed as previously described (Saari et al., 2022 (25) using the High-Capacity cDNA Reverse Transcription Kit (Applied Biosystems; Thermo Fisher Scientific, Waltham, MA, USA) and an Eppendorf MasterCycler thermal cycler (Eppendorf, Hamburg, Germany). The maturation status after DE, PDE, or SIEC differentiation was assessed by TaqMan® (Applied Biosystems, Inc., Foster City, CA, USA) gene expression analysis of *NANOG* (Hs02387400_g1) and *POUF* (OCT4 Hs00999632_g1), *SOX17* (Hs06634937_s1), *FOXA2* (Hs00936490_m1) and *SLC15A* (*PEPT1* (Hs00192639_m1). *GAPDH* (Hs99999905_m1) served as the endogenous control. Reactions were run at least in triplicate, including no-template controls, on a 7300 Real-Time PCR system using standard cycling conditions (2 min at 50 °C, 10 min at 95 °C; 40 cycles of 15 s at 95 °C; finally 1 min at 60 °C). Data were analyzed with SDS software v2.4, and relative expression levels were calculated using Ct values and the 2 ΔΔCt method (42).

#### 2.6.3. RNA sequencing

RNA sequencing was performed essentially as described previously (Freitag et al., 2020; Macosko et al., 2015; Vuorela et al., 2021 (43–45)). Briefly, cDNA synthesis was initiated from 10 ng of RNA using poly(A) -apturing indexing oligos (Integrated DNA Technologies, S1 Table) and a template-switching reverse transcription reaction (10 mM dNTPs (Meridian Bioscience), RiboLock RNase Inhibitor (Thermo Fisher), Maxima H-RTase (Thermo Fisher), 50 μM of template switch oligo (Integrated DNA Technologies), 5 M betaine (Sigma), RNase-free water (Lonza) 30 min RT, 90 min 52 °C). Following exonuclease I (Thermo Scientific) treatment (45 min 37 °C; 85 °C 15 min) to remove unused oligos, cDNA was amplified by PCR (Kapa HiFi HotStart Readymix (Roche), SMART PCR primer (Integrated DNA Technologies), nuclease-free water, and thermocycled in T100 thermocycler (BioRad ; 95°C 3 min; 4x(98 °C 20 sec;65 °C for 45 sec; 72 °C 3 min); 12x (98 °C 20 sec; 67 °C 20 sec; 72°C 3 min); finally 5 min 72 °C), pooled in sets of 48 samples with unique indexes (Supplemental table 3), and purified using magnetic bead–based cleanup (MAGBIO). The 3′-end sequencing libraries were prepared from 1 ng of amplified cDNA using the Nextera XT DNA Library Prep Kit (Illumina), with modified index primers (Supplemental data 3). Libraries were purified, quantified using Qubit (ThermoFisher Scientific), quality-controlled, and stored at -80 °C. Sequencing was performed on an Illumina NextSeq 500 platform using custom primers, generating 20-bp read 1 and 50-bp paired-end read 2. Library preparation and sequencing were carried out at the Biomedicum Functional Genomics Unit (University of Helsinki).

#### 2.6.4 Read alignment and generation of digital expression data

Read alignment and digital gene expression quantification were performed at the Biomedicum Functional Genomics Unit (University of Helsinki). Raw sequencing data quality was assessed using FastQC and MultiQC (46). Reads shorter than 20 bp or of low quality were removed with Trimmomatic (47). Filtered reads were processed using the Drop-seq tools pipeline (45) , including conversion to sorted BAM files, tagging with sample barcodes and unique molecular identifiers (UMIs), and trimming of adapter and poly(A) sequences. Reads were aligned to the GRCh38.p14 genome using STAR (48) with GENCODE release 44 annotations (49). Following alignment, barcode and UMI information was restored, annotation tags were added, and systematic barcode synthesis errors were detected and corrected using Drop-seq tools.

#### 2.6.5 Differential expression analysis

Differential expression analysis was performed with DESeq2 (50) v1.40.2 in R v4.3.0, with stimulation and differentiation samples analyzed separately. Samples with fewer than 100,000 reads were excluded from the analysis. Technical replicates were collapsed with the collapseReplicates function. Wald statistics were calculated using a t-distribution to accommodate smaller sample groups and to provide more conservative p-values. The minimum expected mean was set to 1e-6 to retain possible lower counts per gene or sample from drop-seq. Automatic outlier replacement was disabled to preserve original count values and maintain transparency for sample groups with fewer replicates. To mitigate potential noise from low-expression genes retained after differential expression analysis, we excluded genes with fewer than 50 reads in any stimulation group, as well as those with fewer than 10 reads in any differentiation group, from further analysis. For each scatterplot, the 50 most significant genes by adjusted p-value from each compared contrast were combined, and the log2 fold change of selected genes was plotted with ggplot2 (4.0.1) and ggrepel (0.9.8) in R.

#### 2.6.6 Pathway enrichment

Overrepresentation analysis (ORA) was performed using the enrichGO function from the ClusterProfiler v4.14.6 package (51). Differentially expressed genes were converted to Entrez IDs with the bitr function, using annotations from the org.Hs.eg.db database v3.20.0. Enrichments were assessed across all three Gene Ontology (GO) domains: Biological Process (BP), Molecular Function (MF), and Cellular Component (CC). The p-values were adjusted for multiple testing using the Benjamini-Hochberg (BH) method, with an adjusted p-value threshold of 0.05. Redundant GO terms were merged with the simplify function. Gene Set Enrichment Analysis (GSEA) was conducted using the gseGO function, focusing on BP terms. Genes were ranked by the log2 fold change obtained from differential expression analysis.

#### 2.6.7 Cell type analysis

To assess potential differentiation, normalized expression values from all samples analyzed together were obtained using DESeq2. A curated set of intestinal cell type–specific marker genes was collected from published studies (52–56). These marker genes were used to compare the gene expression values of SIEC CeD and PDE CeD along with their respective controls. Based on the expression levels of these markers, we inferred whether differentiation had occurred and which cell types were most likely represented.

Cell-type deconvolution was performed using CIBERSORTx (57). As a reference, we used Pan-GI Cell Atlas single-cell RNA-seq dataset (58).The dataset was restricted to small intestine and containing only adult donors. Level 2 annotations were used as cell type labels, and cell types with fewer than 50 cells were removed. Cells were downsampled to 1000 cells and 500 genes per cell type. Markers were identified with scanpy (59), and genes expressed in more than 50 % of cells outside their respective cluster and genes with zero expression across samples were excluded.

### 2.7. Functionality assays

#### 2.7.1. Dipeptide uptake assay

SLC15A1/PEPT1 activity was assessed as previously described (Hilgendorf et al Kabeya et al., 2018; Saari et al., 2022 (60–62)). Cells were preincubated in Hank’s balanced salt solution (HBSS, Thermo Fisher, Gibco) for 1 h, followed by incubation with the fluorescent PEPT1 substrate β-Ala-Lys-AMCA (25 μM) in HBSS for 4 h. After uptake, cells were washed with DPBS, fixed in 4% paraformaldehyde, and mounted with Vectashield containing DAPI after excision of insert membranes.

Confocal images were acquired from the apical side using an LSM780 confocal microscope (Zeiss) with a Plan-Apochromat 40x/1.4 NA oil objective. DAPI and AMCA fluorescence were excited at 405 nm and 488 nm, respectively, and acquired sequentially using GaAsP-PMT detectors. Z-stacks were collected without averaging using a 1 Airy unit pinhole, 0.50 μs dwell time, 99 nm xy pixel size, and 1 μm z-steps, starting above the apical surface and extending to the lowest DAPI-positive plane.

Images were converted to TIFF format, channels separated, and z-stacks processed in Fiji ImageJ. Background (negative control) was defined from planes above the cell layer, while PEPT1 signal was quantified from planes corresponding to the nuclear midpoint. Final images were assembled in Adobe Photoshop (2026).

#### 2.7.2. Cytotoxicity assay

Cytokine stimulations are known to damage cells and thus increase the release of lactate dehydrogenase (LDH) into cell culture media (37). Thus, the cytotoxic effect of the treatments was evaluated by measuring the LDH activity of the culture media with the Cytotoxicity detection kit (Roche Diagnostics GmbH, Mannheim, DE) according to the manufacturer’s protocol. The result was read with a VICTOR® Nivo™ multiplate reader (Perkin Elmer, Shelton, Connecticut, USA) with a preset program of 490 nm 0.1s, and 650 nm 0.1s. The readings were correlated to the untreated sample.

### 2.8. Statistical significance

The statistical significance of numerical data was analyzed using IBM SPSS Statistics version 26, with a two-tailed Mann–Whitney U test. The number of biological and technical replicates is indicated in the figure legends.

### 2.9. Ethical Issues

The study was approved by the Ethics Committee of Pirkanmaa Hospital District to derive and expand iPSC lines (R08070 and R12123), and written consent was obtained from all fibroblast donors. All patients were over 18 years of age. The use of cell lines for gastrointestinal research was approved by the Ethical Committee of Pirkanmaa Hospital District (R20008). No new cell lines were derived in this study.

## 3. Results

### 3.1. All cell lines had the ability to differentiate towards SIECs

Three of the used iPS cell lines were new. For that reason, the cell differentiation was evaluated using wide-field microscopy, RT-PCR, and RNA sequencing. At the iPSC stage, all lines expressed pluripotency markers *NANOG* and *POU5F1* at both RNA and protein levels (Fig 1A; S1 Fig A, B). After three days, definitive endoderm (DE) markers *FOXA2* and *SOX17* were detected in all lines (Fig 1B, S1 Fig C), and by day 7, posterior definitive endoderm (PDE) markers *CDX2* and SOX17 were expressed (Fig 1C, S1 Fig D).

**Fig 1.**
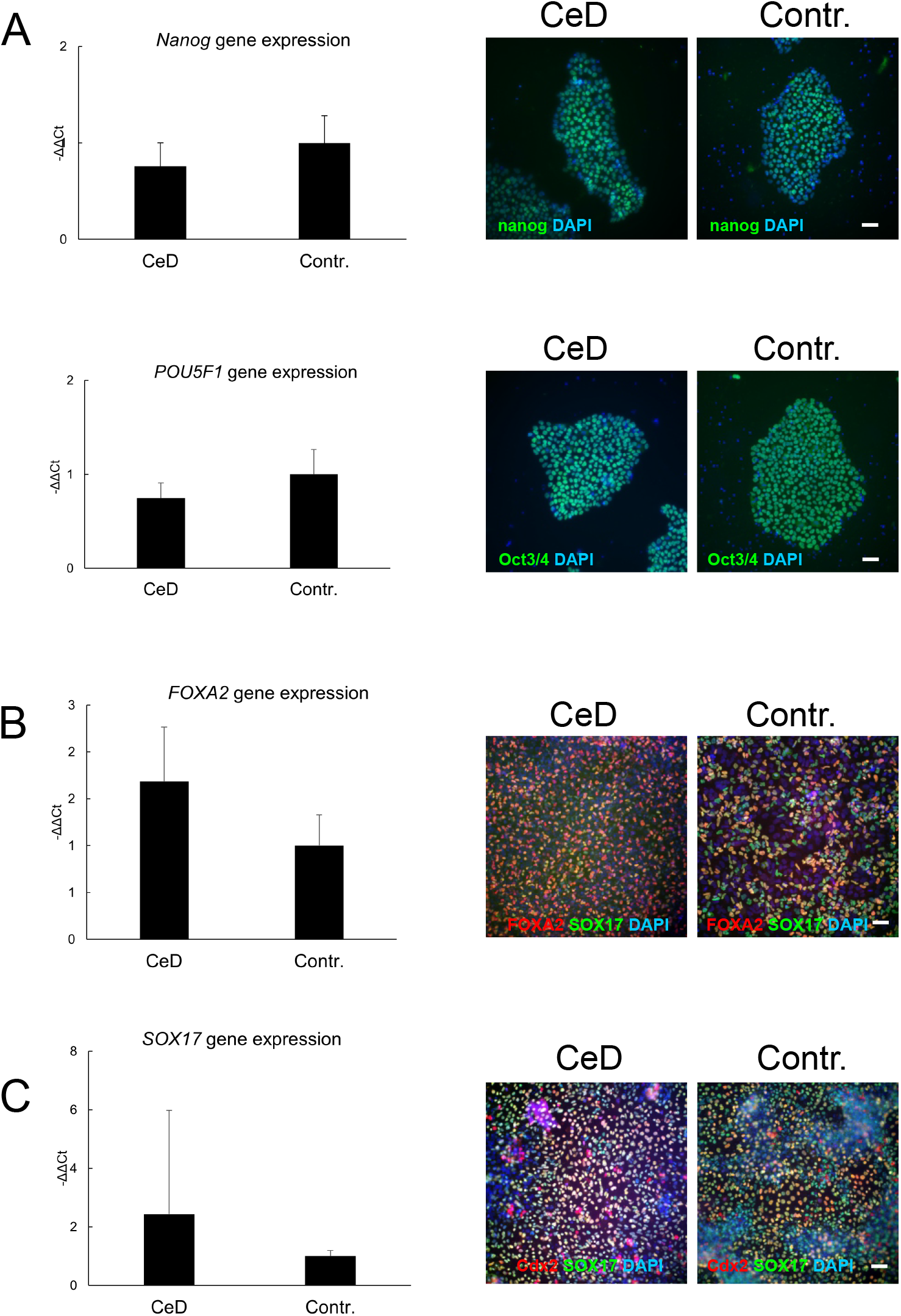
Differentiation of the CeD-derived cell line and the healthy-control-derived cell line iPSCs towards DE, PDE, and SIEC. Before differentiation, both iPSC lines expressed the as well as the *Nanog* gene and protein (n = 3), and *POU5F1* / Oct3/4 gene and protein (n = 3) (panel A). At the DE stage, cultures expressed the *FOXA2* and *SOX17* gene and protein (n = 3; panel B, FOXA2 red, SOX17 green, DAPI blue, 100 μm). At the PDE stage, cultures expressed the *Cdx2* and *SOX17* gene and protein (n = 3; panel C, Cdx2 red, SOX17 green, DAPI blue, scale bar 100 μm). Microscope images (A-C) were taken with an Evident-Olympus IX83 microscope 20x lens using a Hamamatsu ORCA-Fusion C14440-20UP camera and NEO LiveImaging software.

To visualize cellular morphology and culture organization, representative images of a single Transwell culture are shown in Fig. 2A. Phase-contrast microscopy indicated a homogeneous epithelial morphology, and confocal microscopy confirmed the formation of an EpCAM- and PEPT1-positive epithelial layer. Elongated phalloidin- and DAPI-positive cells were detected beneath the epithelial compartment. Despite the presence of a morphologically uniform and polarized epithelial monolayer, TEER measurements were relatively low, varying between 20 and 840 Ωcm^2^; the culture depicted in Fig. 2A exhibited a TEER value of 108 Ωcm^2^. The iPSC-SIECs exhibited a polarized epithelial phenotype, characterized by E-cadherin localization at lateral cell junctions adjacent to the apical membrane and villin-positive brush-border structures, indicative of enterocyte maturation (Fig 2B). TEER of the CeD- derived patient culture in the Fig 2B was 22 Ωcm2, and healthy control derived culture in the Fig 2B 20 Ωcm^2^. At day 28, iPSC SIECs expressed *SLC15A1* mRNA and PEPT1 protein (Fig 6A), and uptake of a fluorescent dipeptide substrate confirmed PEPT1 activity (Fig 2C; S1).

**Fig. 2.**
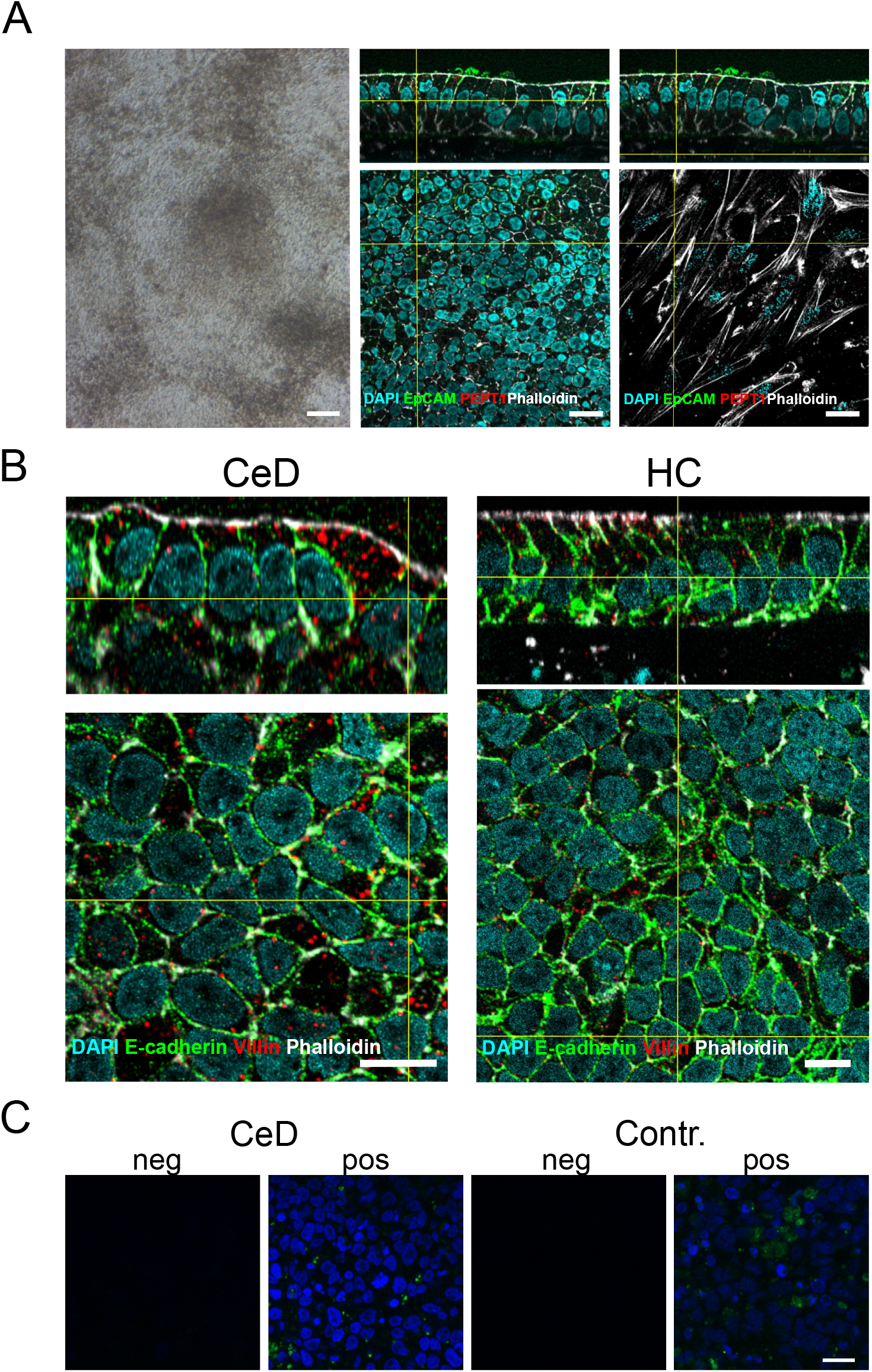
Differentiation of celiac disease (CeD)-derived line and healthy control-derived line iPSCs into small intestinal epithelial cell (SIEC) cultures. (A) Representative cell culture morphology and its relationship to transepithelial electrical resistance (TEER) (CeD n = 1, control n = 1; technical replicates: CeD n = 18, control n = 9). The left panel shows a phase-contrast image of a day-25 culture with a TEER value of 108 Ω·cm^2^ (scale bar, 100 μm). The middle and right panels show confocal images of the same culture. The middle panel depicts an EpCAM-positive (green) and PEPT1-positive (red) epithelial layer, whereas the right panel shows the underlying layer containing elongated cells stained only with DAPI (blue) and phalloidin (white) (scale bars 10 μm). Phase-contrast images were acquired using a Zeiss Axiovert A1 microscope equipped with a 5x LD A-Plan objective and a Zeiss Axiocam ERc 5s camera. (B) After 28 days of differentiation, iPSC-SIECs expressed the adherens junction marker E-cadherin (green) and the brush-border marker villin (red), together with phalloidin (white) and DAPI (cyan). Confocal images in panels A and B were acquired using an Evident FLUOVIEW FV4000 laser-scanning confocal microscope equipped with a 60x silicone oil immersion objective. Sample numbers were as follows: villin staining, CeD n = 2 and control n = 1; E-cadherin staining, CeD n = 1 and control n = 1. Scale bars, 10 μm. (C) Intestinal epithelial cell specific functionality assessed with PEPT1 dipeptide uptake assay. On left there is image captured above the cellular level and on the right, there is image taken from the level which is in the middle of nucleus (CeD n=3; Contr. n=3; dipeptide D-Ala-Leu-Lys- AMCA green, DAPI blue, 20 μm). Confocal images were taken with a Zeiss LSM780 laser scanning microscope with a 40x lens and ZEN Blue 3.6 software.

Cross differentiation stages RNA sequencing revealed gradual transcriptomic changes. Among the most significantly altered genes, the enterocyte-associated APOA1 expression increased from the PDE stage onward (Fig 3A B; S2 Fig B). APOA1 and DPP4 in CeD-derived line had higher expression in PDE-stage than in SIEC-stage, whereas in healthy control-derived line both genes had higher expression in SIEC- than in PDE-stage (Fig 3B). TTR gene increased its expression CeD-derived line and healthy control-derived cell line in SIEC-stage (Fig 3A). Similar differentiation associated expression patterns were observed in T1D-derived line and T2D-derived line (S3 Fig A).

**Fig 3.**
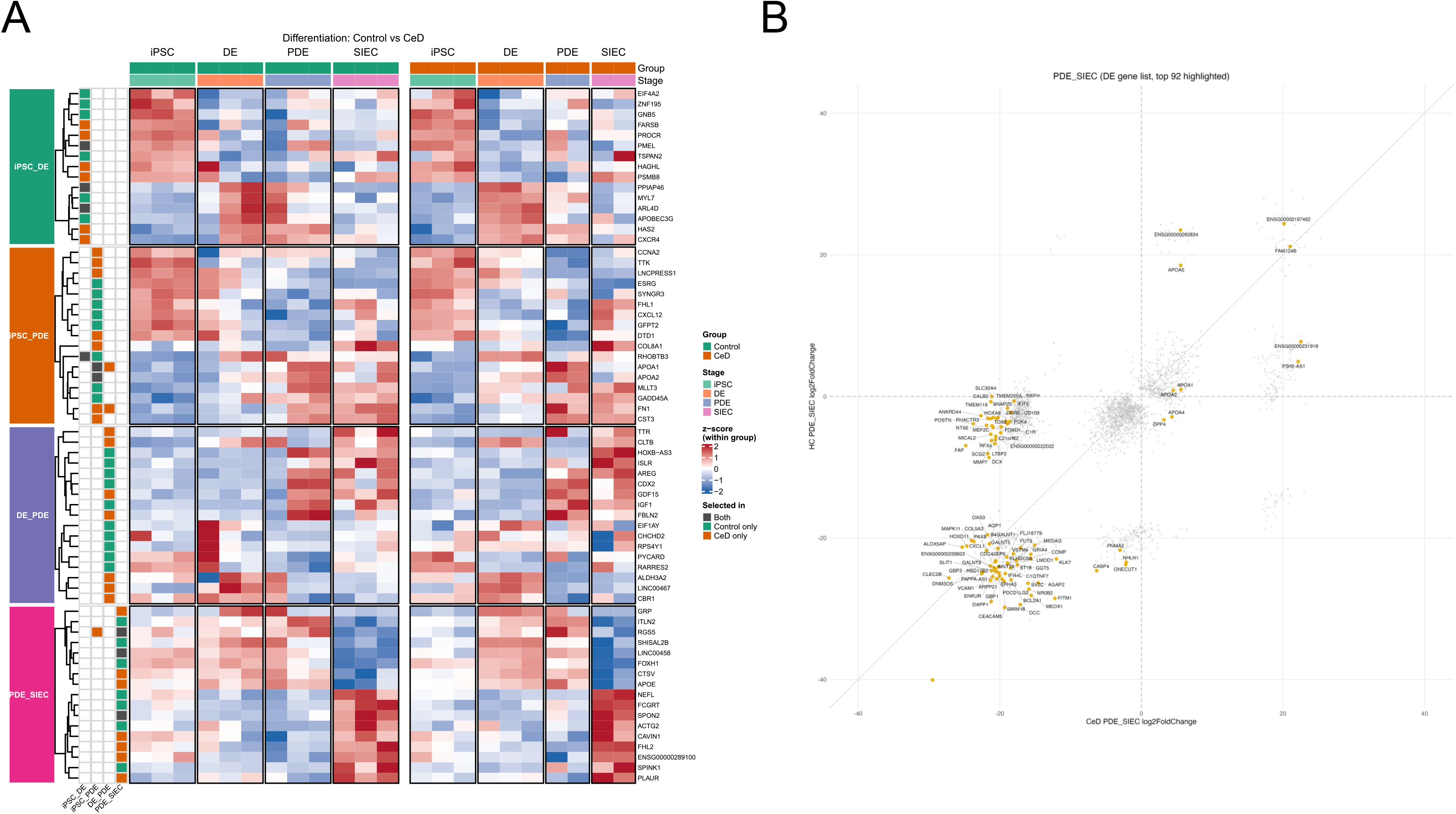
Heatmap of RNA-seq data including differentially expressed genes during each stage of differentiation. The mRNAs collected from (A) CeD-derived cell line and healthy-control-derived cell line l (Contr.) at the iPSC, DE, PDE, and SIEC stage were analyzed. Analysis was achieved from iPSC (CeD n = 3; Contr. n = 3); DE (CeD n = 1, Contr. n = 2); PDE (CeD n = 2, Contr. n = 3) and SIEC (CeD n = 2, Contr. n = 3). The RNA sequencing and results were separately plotted into heatmaps of the 50 genes which exhibited statistically most significant expression change. (B) Scatter plots of differentially expressed genes (log2) in PDE- vs SIEC-stage between the CeD-derived cell line and the healthy-control- derived cell line.

Analysis of shared differentially expressed genes in Venn diagrams showed that the CeD-derived line exhibited fewer transcriptional changes between iPSC and DE stages compared with the healthy- control-derived line, but a greater overlap between the iPSC-DE and DE–PDE transitions; the PDE-SIEC transition was comparable between groups (Fig 4A, BS3). Gene Ontology and Gene Set Enrichment Analyses demonstrated stage-specific pathway activation: iPSC-DE transition was enriched for anterior– posterior patterning and tissue morphogenesis, DE–PDE for embryonic organ and tissue development, and PDE–SIEC for extracellular matrix organization, ribosomal biogenesis, and epithelial maturation (Fig 4C-D).

**Fig 4.**
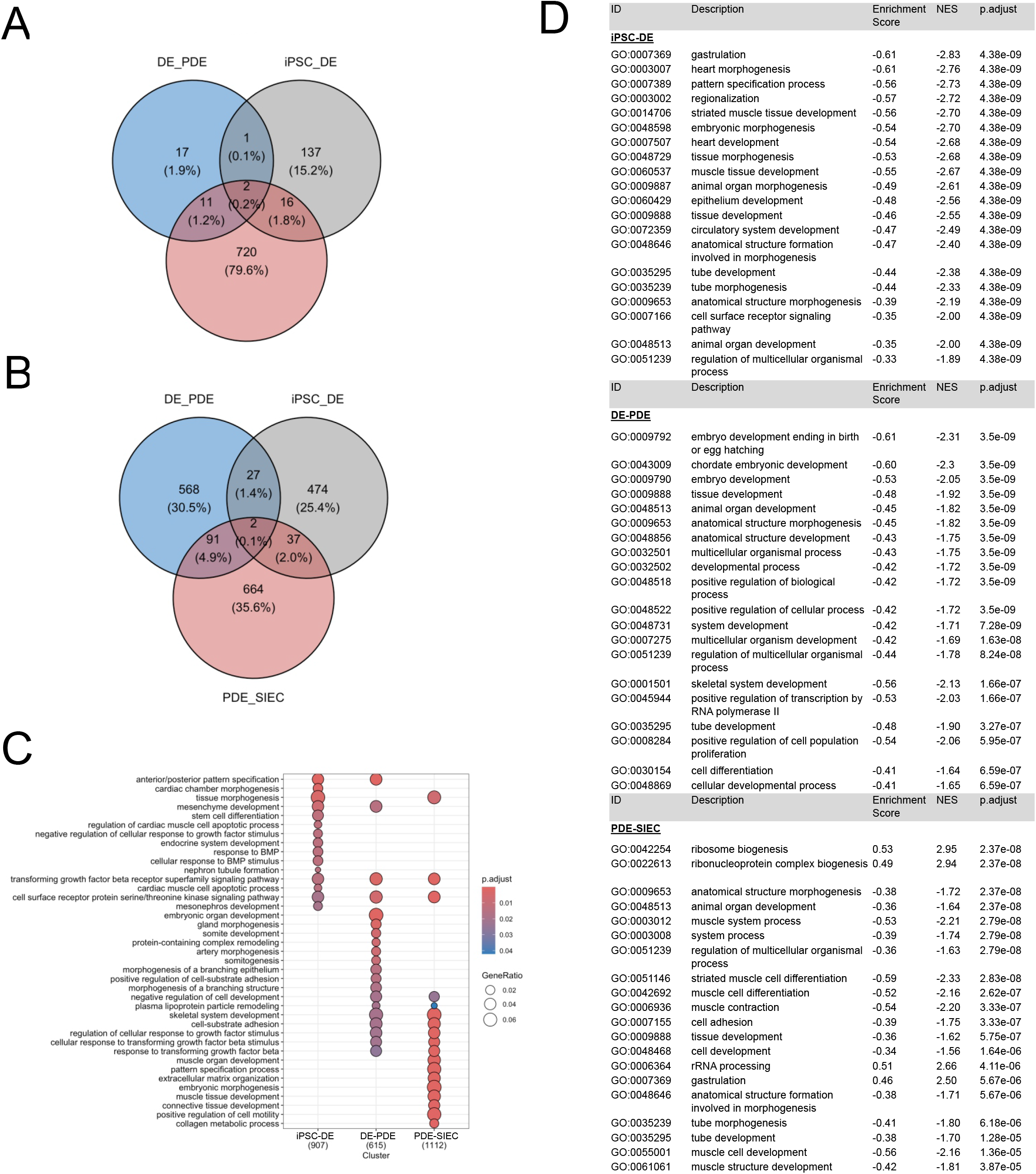
Venn diagram of shared genes and Gene Ontology expression analysis of the CeD-derived cell line and the healthy-control-derived cell line. A) Venn diagrams illustrating the genes showing the most significant expression changes identified by RNA sequencing and shared across developmental stages in CeD**-**derived cell line (A) and in healthy-control-derived cell line (B) samples; Gene Ontology (GO) enrichment analysis (C) of all significantly changed biological processes across developmental stages—iPSCs vs. DE (grey), DE vs. PDE (blue), and PDE vs. SIECs (red)—in CeD and control samples. (D) Gene set enrichment analysis (GSEA) of the 20 most significantly changed gene classes in pairwise comparisons (iPSCs vs. DE, DE vs. PDE, PDE vs. SIECs) in CeD-derived line, healthy control-derived line, T1D-derived line, and T2D-derived line. Normalized enrichment score (NES) and adjusted p-value (p-adjust).

Gene set enrichment analysis (GSEA) of the CeD-derived line and the healthy-control-derived line T1D-derived line, and T2D-derived line revealed stage-specific pathway enrichment during differentiation. At the DE stage, gene sets related to gastrulation, pattern specification, regionalization, and tissue morphogenesis were enriched. During the DE–PDE transition, enrichment was observed in pathways associated with embryo differentiation, tissue development, and anatomical structure morphogenesis. In the PDE–SIEC transition, gene sets involved in ribosome and ribonucleoprotein complex biogenesis showed positive enrichment, while pathways related to cell and tissue development and morphogenesis were upregulated in SIECs compared with PDEs (Fig 4C-D, S4).

Bioinformatic cell type composition analysis was used to characterize differentiation-associated cell populations based on established marker genes (52–56). Both cell lines showed strong activation of transit-amplifying cell markers and progressive upregulation of immature and mature enterocyte genes at the SIEC stage (Fig 4A). After the PDE stage, transcriptional signatures of multiple intestinal epithelial lineages including M cells, Paneth, goblet, and enteroendocrine cells emerged. In addition, myofibroblast-associated genes were induced, with stronger activation in CeD-derived cells than in controls (Fig 5A).

**Fig 5.**
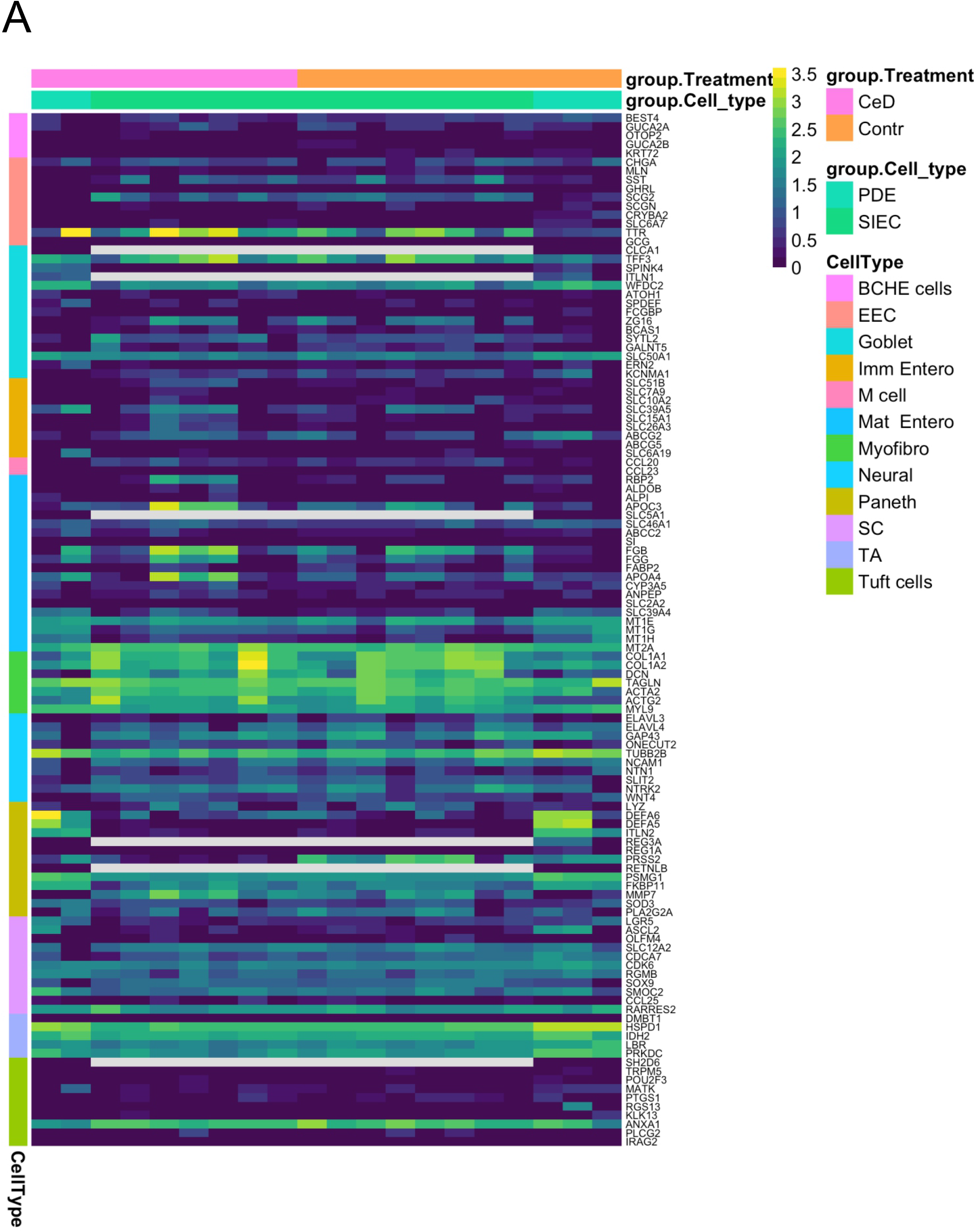
Cell type composition analysis and immune reaction–related genes within differentially expressed genes. (A) cell type composition analysis performed by visualizing our gene expression data from reference gene sets corresponding to different cell types (A) (52–56) in the CeD-derived line (PDE n = 2, SIEC n = 7) and the healthy-control-derived line CeD (PDE n = 3, SIEC n = 8) samples. Abbreviations: Best4+ cells (BCHE), enteroendocrine cell (EEC), Goblet cell (Goblet), immature enterocyte (ImmEntero), immature enterocyte (Imm Entero), microfold cell (M cell), mature enterocyte (Mat Entero), myofibroblast (Myofibro), late immature enterocyte (LI_entero), stem cell (SC), transit- amplifying cell (TA), Paneth cell (Paneth).

### 3.2. Morphology and gene expression of the iPSC-SIECs after cytokine stimuli

The iPSC-SIECs derived from the CeD-derived line and the healthy-control-derived line, including disease controls, T1D-derived line and T2D-derived line, were stimulated with cytokines IFNγ and TNFα linked to CeD (28–30,55). The dipeptide transporter PEPT1 was expressed at protein level (Fig 6A and S4 Fig A) and at mRNA level (Fig 6B, and S4 Fig B) across all groups.

**Fig 6.**
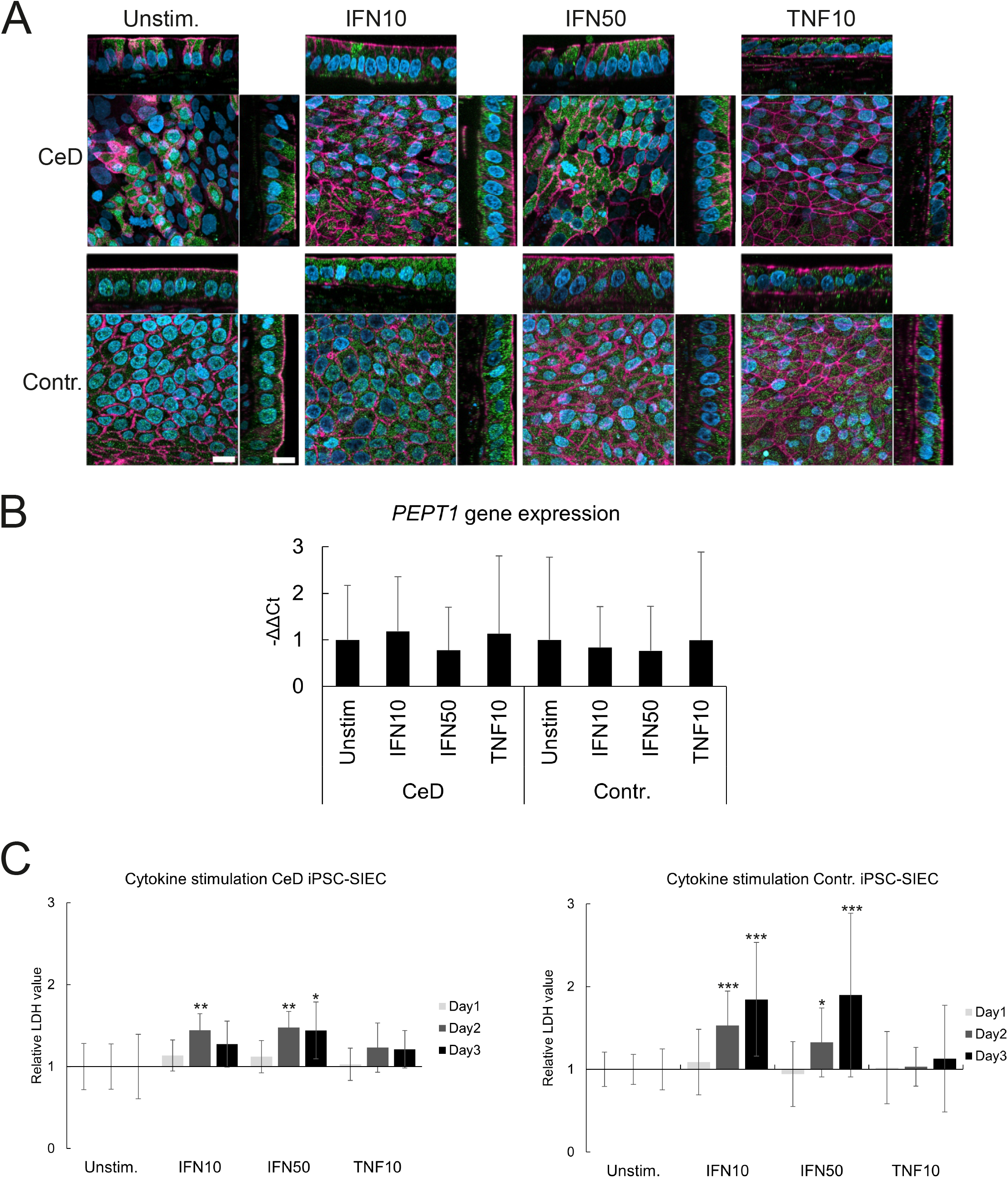
**Cellular morphology and gene expression, as well as cytotoxicity, following cytokine stimuli**. SIECs of the CeD-derived line (CeD) and the healthy-control-derived line (Contr.) iPSCs and matured for 26 days were stimulated with IFNγ (10 ng/mL and 50 ng/mL) or TNFα (10 ng/mL) for 48 h. A) Immunofluorescence staining for PEPT1 (green), filamentous actin (phalloidin, red), and nuclei (DAPI, blue). B) PEPT1 expression quantified by qRT-PCR (CeD n = 6, Contr. n = 3). Confocal images were taken with a Zeiss LSM800 laser scanning microscope with a 63x lens and ZEN Blue 3.6 software. C) Cytotoxicity assessed based on lactate dehydrogenase (LDH) activity in culture media after 48 h of cytokine exposure compared to unstimulated controls (n = 3). Statistical significance * = p ≤ 0.05; * = p ≤ 0.005, and *** = p ≤ 0.0005.

Cytokine effects were evaluated by measuring lactate dehydrogenase (LDH) release, which was significantly increased by IFNγ in both the CeD-derived line and the healthy-control-derived line iPSC- SIECs (Fig 6C; S4 Fig C).

### 3.3. Cytokines induced gene expression alterations

The effects of 48-h cytokine stimulation were assessed by RNA sequencing. IFNγ at both 10 and 50 ng/mL robustly induced interferon responsive pathways, including strong upregulation of IRF family members (*IRF1, IFIT2, IFIT3, IDO1*), chemokines (*CXCL9, CXCL10, CXCL11*), HLA class I-genes (*HLA-A, -B, -C, -E, -F*), and additional interferon-associated genes such as *STAT1, WARS1, IL15*, and *PSMB9* (Fig 7A;S5A-D). Transcriptional responses were broadly similar across the CeD-derived line and the healthy-control-derived line (Fig 7B-C) iPSC-SIECs at both IFNγ concentrations. Although there were minor differences: at 10 ng/mL IFNγ 93 genes were differentially expressed in the CeD-derived line versus 283 in the healthy-control-derived line (Fig 7B), and at 50 ng/mL 145 genes in the CeD-derived cell line versus 331 in healthy control-derived cell line (Fig 7C). Slightly lower induction in the CeD-derived cell line was observed for some interferon-responsive genes, including *HLA-B, HLA-C, ICAM1, IRF1,* STAT1, *TAP1/2, PSMB8/9*, and *WARS1* (Fig 7B-C).

**Fig 7.**
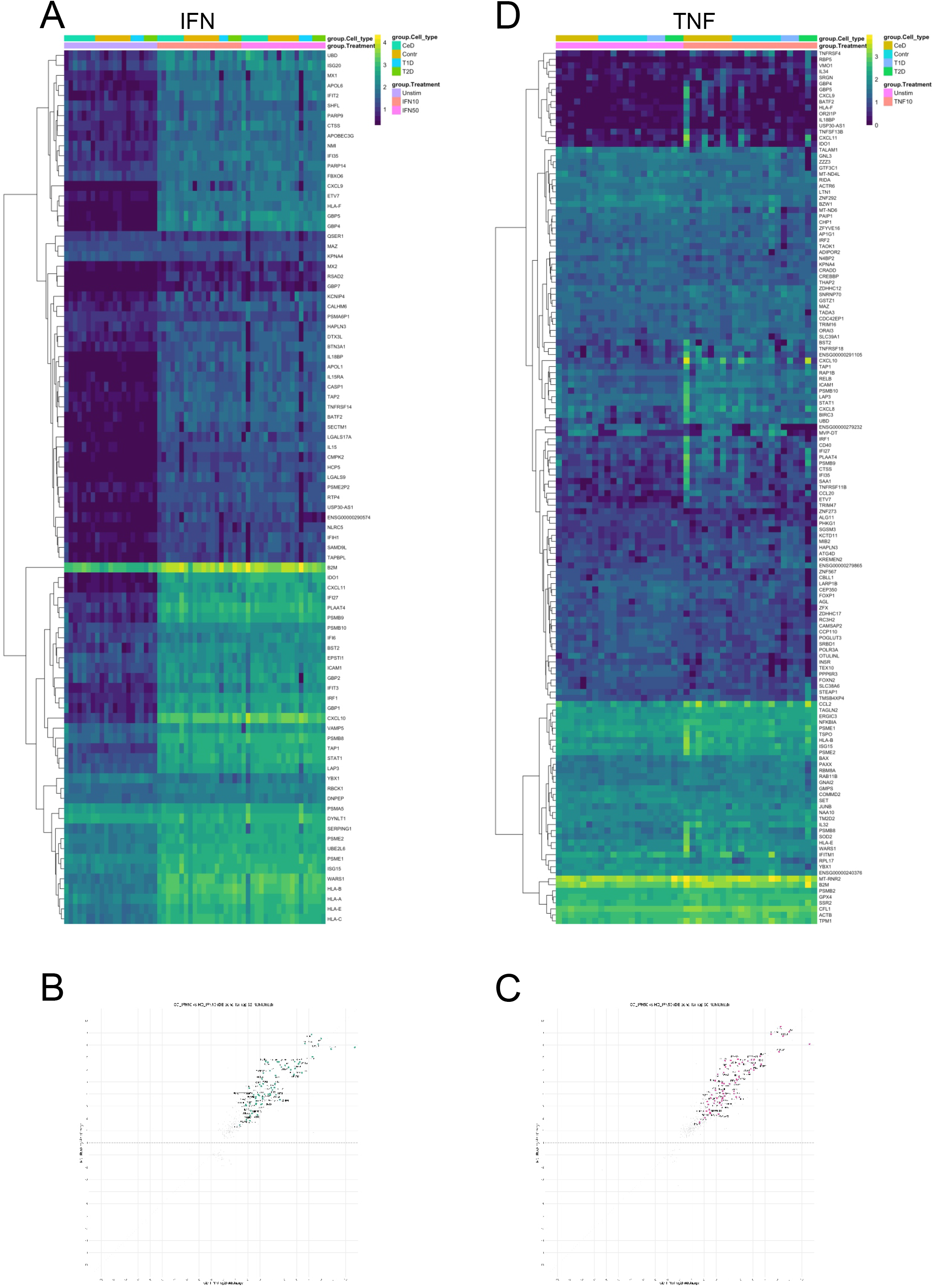
Heatmaps and scatter plots of significantly differentially expressed genes in cytokine-stimulated PSC-SIECs. Transcriptomic responses to cytokine stimulation in iPSC-derived SIECs. The CeD-derived ine (CeD), and the healthy-control-derived line (Contr.), type 1 diabetes -derived cell line (T1D), and type 2 diabetes -derived cell line (T2D) iPSC-SIECs were treated with IFNγ 10 ng/mL (A, B) or 50 ng/ ml (A, C), or TNFα (10 ng/mL) (D) for 48 h. Gene expression changes were analyzed with RNA sequencing and visualized as heatmaps and scatter plots (CeD n = 5, Contr. n= 7, T1D n= 3, T2D n = 3).

TNFα stimulation elicited more variable transcriptional responses. At 10 ng/mL, TNFα significantly upregulated a limited set of shared genes in the CeD-derived line and the healthy-control-derived line iPSC-SIECs, including *CXCL8, CXCL10, CXCL11, CCL2, ICAM1*, and *UBD* (Fig 7B, S6 Fig A-C). No apoptotic pathway activation was detected, as caspase expression remained unchanged, while *NFKBIA* was significantly upregulated in all (CeD-derived line, and the healthy-control-derived line, type 1 diabetes -derived cell line and type 2 diabetes -derived cell line) iPSC-SIECs. The TNFα induced strong upregulation of several proteasome-related genes (*PSME1, PSME2, PSMB2, PSMB10*) specifically in CeD-derived line (Fig 7C, S6 A-D).

### 3.4. Gene Ontology processes induced by cytokine stimulation

To assess the impact of IFNγ stimulation, differentially expressed genes were grouped by Gene Ontology (GO) biological processes. Treatment with 10 ng/mL IFNγ for 48 h resulted in strong enrichment across all cell lines for antiviral and immune related processes, including response to virus, defense response to virus, and innate immune response, as well as regulation of immune responses (Fig 8A).

**Fig 8.**
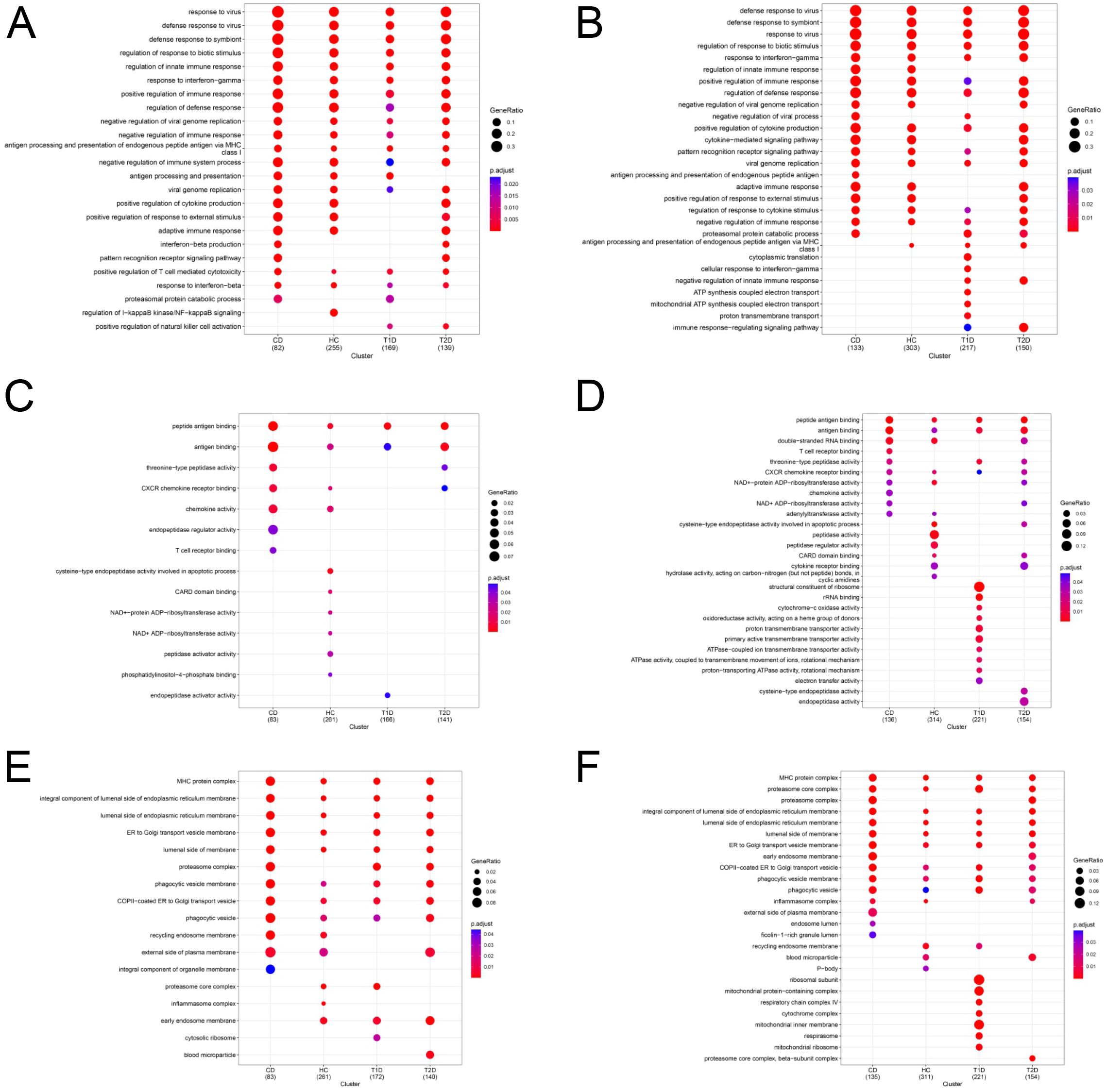
**Bubble chart of biological processes (A, B), molecular processes (C, D), and cellular components (E, F) induced by IFN**γ **cytokine stimuli.** The samples in A, C, and E were stimulated with 10 ng/ml of IFNγ and those in B, D, and F with 50 ng/ml of IFNγ. Gene expression data from the CeD- derived line (CeD), type 1 diabetic -derived cell line (T1D), type 2 diabetic -derived cell line (T2D), and the healthy-control-derived cell line healthy control (Contr.) lines matured for 26 days.

iPSC-SIECs of the CeD-derived line and the healthy-control-derived line, and the T2D -derive line showed significant enrichment of pathways related to positive regulation of cytokine production, adaptive immune response, and response to external stimuli, whereas these processes were not significantly induced in T1D-derived cell line. Consistently, the adaptive immune system pathway was enriched in all cell types except T1D-derived cell line. Stimulation with 50 ng/mL IFNγ largely recapitulated the effects of 10 ng/mL, with comparable pathways, gene ratios, and statistical significance, indicating a dose- consistent response.

At the molecular processes, antigen binding and peptide antigen binding were consistently enriched across all (CeD-derived line, and the healthy-control-derived line, type 1 diabetes -derived cell line and type 2 diabetes -derived cell line) cell lines at both IFNγ concentrations, while CXCR chemokine receptor binding was additionally induced at 50 ng/ml. IFNγ also altered multiple cellular component categories, including MHC protein complexes, ER and ER-to-Golgi transport vesicles, and phagocytic vesicle membranes, with similar enrichment patterns across cell lines and doses (Fig 8C-D).

In contrast, TNFα stimulation (10 ng/mL) elicited limited and heterogeneous responses. From the biological processes the response to chemokine was shared between the CeD-derived line and the healthy-control-derived line, and T2D-derived line. No other common molecular function or cellular component pathways were observed across all four cell lines.

### 4.1. Small intestinal cell deconvolution for cytokine stimulated samples

Cellular responses to inflammatory stimuli can vary between cell types, and mesenchymal and neural cell populations have previously been reported to exhibit greater responsiveness to IFNγ stimulation (54). Thus, ellular composition in CeD-derived line, healthy control-derived line, T1D-derived line, and T2D-derived ine iPSC-SIECs were evaluated using CIBERSORTx cell-type deconvolution analysis. Under unstimulated onditions, fibroblasts comprised 4.8%-7.8% of cells, myofibroblasts 3.9%-5.3%, and smooth muscle cells 1.2% (Fig. 9A). Following treatment with 10 ng/ml IFNγ, the T1D-derived line exhibited a lower fibroblast proportion (3.9%) than the CeD-derived (6.9%), healthy control-derived line (7.0%), and T2D-derived line 7.8%), whereas myofibroblast (2.8%-4.0%) and smooth muscle cell (0.8%-1.2%) proportions remained omparable across all lines (Fig. 9B). A similar pattern was observed following 50 ng/ml IFNγ treatment, with fibroblasts accounting for 4.0% of cells in the T1D-derived line and 6.7%-7.5% in the other lines. In ontrast, myofibroblasts were slightly more abundant in the T1D-derived line (5.4%) than in the remaining ines (2.8%-3.9%), while smooth muscle cell proportions remained consistent (0.4%-0.7%) (Fig. 9B).

**Fig 9.**
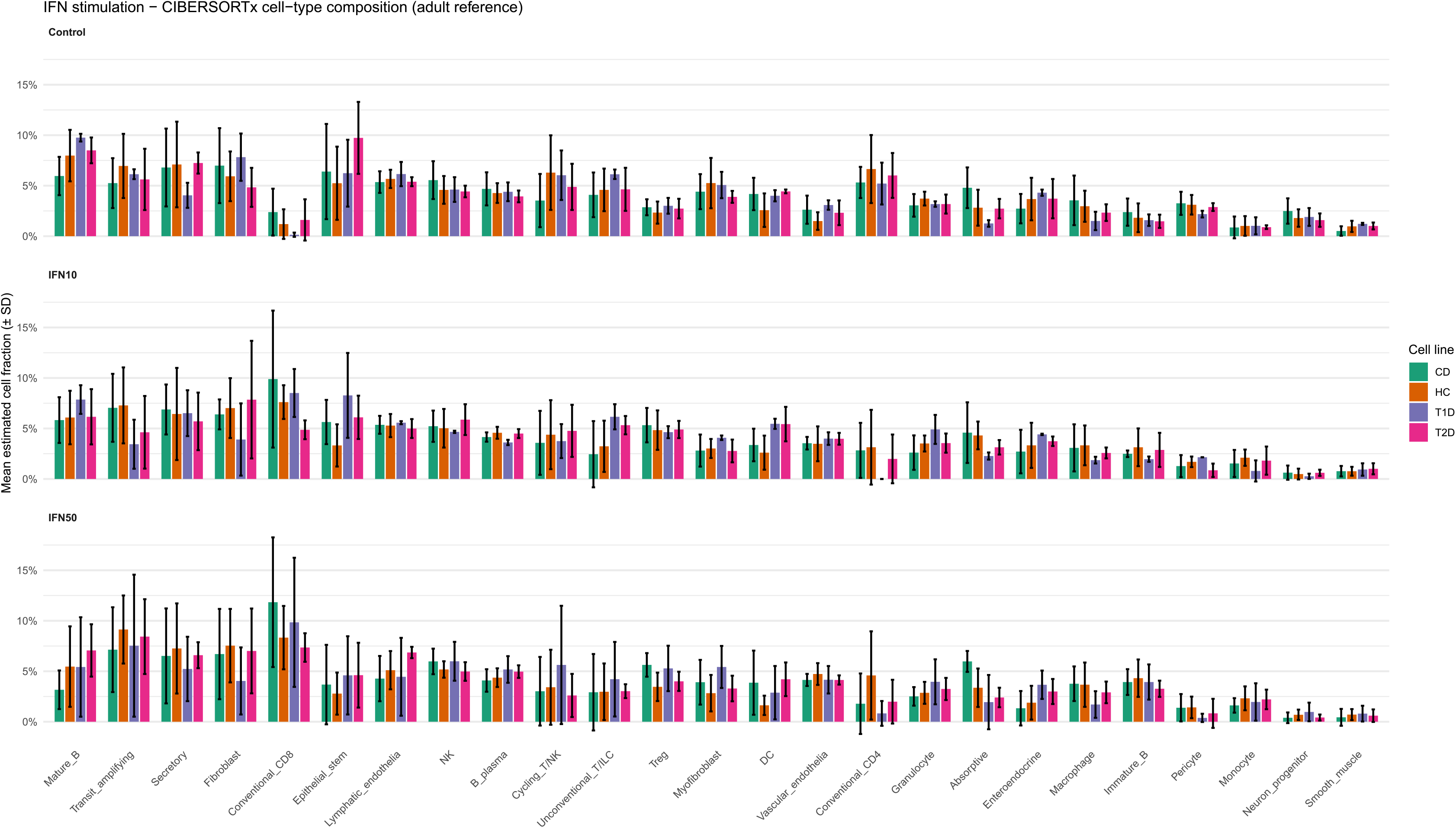
Cell-type deconvolution with CIBERSORTx. using adult small intestinal data (58) to assess proportions of different intestinal cell type**s** in unstimulated, 10 ng/ml of IFNγ and 50 ng/ ml 50 ng/ml of IFNγ stimulated iPSC-SIECs from the CeD-derived line (CeD), healthy-control-derived cell line, type-1- diabetes -derived cell line (T1D) and type-2-diabetes -derived cell line (T2D). Mature B-cell (Mature_B); Natural Killer (NK), plasma B-cells (B_plasma); cycling T cells/ Natural Killer (Cycling_T/NK), Unvonventional T cells innate lymphoid cell (Unconventional_T/ILC), T regulatory cells (Treg); Dendritic cells (DC), Immature B cells (Immature_B).

The combined proportion of mesenchymal cells (fibroblasts, myofibroblasts, and smooth muscle cells) in he CeD-derived line was 11.9%, 10.0%, and 11.2% in unstimulated, 10 ng/ml and 50 ng/ml IFNγ timulated, respectively. In the healthy control-derived line were 12.2%, 10.8%, and 11.1% in unstimulated, 10 ng/ml and 50 ng/ml IFNγ stimulated, respectively. Overall, the mesenchymal cell populations exhibited only minor variation between cell lines and treatment conditions.

Among epithelial cell populations, epithelial stem cells constituted 5.2%-6.4% of cells in the CeD- derived line, healthy control-derived line, and T1D-derived line, whereas a higher proportion was observed n the T2D-derived line (9.7%). IFNγ treatment reduced epithelial stem cell proportions in a dose-dependent manner in the CeD-derived line, from 6.4% under unstimulated conditions to 5.6% and 3.7% following reatment with 10 ng/ml and 50 ng/ml IFNγ, respectively. A similar decrease was observed in the healthy ontrol-derived line, in which epithelial stem cells declined from 5.6% to 3.3% and 2.8%, respectively. In he T2D-derived line, epithelial stem cell proportions decreased from 9.7% to 6.1% following IFNγ reatment.

Absorptive enterocytes accounted for 1.3%-4.8% of cells, with the lowest proportion observed in the unstimulated T1D-derived line (1.3%) and the highest in the unstimulated CeD-derived line (4.8%). Treatment with 50 ng/ml IFNγ increased the proportion of absorptive enterocytes in the CeD-derived line rom 4.8% to 6.0%, whereas no substantial changes were detected in the other lines. Treatment with 10 ng/mL IFNγ had only minor effect on absorptive enterocyte abundance in the CeD-derived and healthy ontrol-derived lines; however, treatment with 50 ng/ml IFNγ reduced absorptive enterocyte proportions to 1.3% and 1.9%, respectively.

When the proportions of epithelial cell types (epithelial stem cells, transit-amplifying cells, nteroendocrine cells, secretory cells, and absorptive cells) were compared to all cell populations, the CeD- derived cell line contained 26.0%, 26.9%, and 24.6% epithelial cells in unstimulated conditions, after reatment with 10 ng/ml and 50 ng/ml IFNγ, respectively. In healthy control-derived cell line, the orresponding proportions were 25.8%, 24.7%, and 24.4%, respectively.

To assess potential immune priming, immune cell deconvolution analysis was performed on unstimulated PSC-SIECs (Fig 9). Transcripts associated with immune cell populations were detected in both the CeD- derived line and healthy control-derived line, and the expression levels of most immune cell-associated ranscripts were comparable between all four (CeD-derived line, and the healthy-control-derived line, type 1 diabetes -derived cell line and type 2 diabetes -derived cell line) lines. The only differences observed were ower proportion of cycling T cells/natural killer cells and conventional CD4 T cells in the unstimulated CeD-derived line compared with the healthy control-derived line. To identify the genes underlying these mmune cell-associated signatures, the transcripts contributing to the immune cell deconvolution results were subsequently examined in more detail (S7). Analysis revealed that most of immune cell clustered ranscripts are not only immune cell specific genes, but genes, such as *beta-actin* (*ACTB*), *HLA* genes, which re expressed also by other cell types.

## 4. Discussion

Most protocols for differentiating iPSCs into small intestinal epithelial cells (SIECs) rely on intestinal organoid formation in Matrigel (20,21,63), which provides key cues for epithelial polarization (64) but limits access to apical stimuli due to the closed organoid structure. This limitation is commonly addressed by dissociating organoids and re-culturing cells in two-dimensional formats to enable apical and basolateral access (65). Here, our first aim was to bypass the organoid-to-2D transition during iPSC-SIEC differentiation. Our second aim was to compare the differentiation capacity of the CeD-derived line and the healthy-control-derived line iPSCs and to assess their responses to cytokine stimulation.

Differentiation of iPSCs into DE, PDE, and SIECs was verified by stage-specific marker expression. At the DE stage, cultures expressed *SOX17* and *FOXA2*, indicating successful endoderm specification (66,67), accompanied by enrichment of biological processes related to stem cell differentiation, mesenchyme development, and anterior-posterior patterning (68). At the PDE stage, expression of *CDX2* confirmed mid-/hindgut identity (22,23,25,69). At the SIEC stage, iPSC SIECs formed polarized epithelial monolayers and expressed PEPT1 (23,69,70), villin (21,69,71), E-cadherin (21,69), DPP4 (21,71) and EpCAM (56) demonstrating appropriate epithelial organization and enterocyte differentiation. The TEER values of transwell-cultured iPSC-SIECs were lower than those typically reported for Caco-2 cells, consistent with previous reports (72,73), was the same or bit lower than previously reported in transwell cultured iPSC-SIECs (72,74), but similar to that of the human small intestine (73). Cultures with a similar polarized morphology as assessed by confocal microscopy exhibited substantial differences in TEER measurements. The mechanisms underlying this variability are currently unknown and require further investigation.

Gene ontology analysis of the PDE-SIEC transition revealed enrichment of muscle related processes, including muscle system process and muscle cell differentiation, suggesting the presence of mesenchymal cells in iPSC SIEC cultures. This is consistent with evidence that intestinal epithelial and mesenchymal lineages arise from shared developmental trajectories and can co-develop from iPSCs (21,75). In many intestinal differentiation protocols, EpCAM^+^ purification is used before final maturation, although mesenchymal cells have still been detected after purification (22,54,63). Here, EpCAM^+^ selection was intentionally omitted, and differentiation was performed directly on a planar surface without intermediate detachment or replating. Consequently, it remains unclear whether mesenchymal cells emerged during PDE differentiation or later during SIEC maturation. Mesenchymal cells and their secreted factors have been suggested to support long-term maintenance of iPSC-derived epithelial cells (54). Consistent with this, the epithelial cells in iPSC-SIEC cultures that contained mesenchymal cells were highly polarized.

Previous studies have shown that 2D cultures derived from small intestinal biopsies express higher levels of drug transporters than 2D iPSC-derived intestinal organoids, although the latter contain additional cell types, including neuronal, endothelial, and mesenchymal cells (76). In addition the iPSC-SIECs cultured in intestine-on-chip are shown to express higher amounts of *CYP3A4* than transwell or organoid cultured iPSC-SIECs (63) In our system, expression of the enterocyte maturation marker *CYP3A4* was comparable to earlier protocols from the Mizuguchi laboratory (23), but lower than levels achieved with their most recent optimized method (69). No direct comparison with other intestinal cell models was performed in this study, but in our previous study, the same healthy control patient -derived line iPSC-SIECs exhibited higher expression levels of the *CYP3A4* than Caco-2 cells (62).

Both the CeD- and the healthy-control-derived line iPSC-SIECs expressed a broad range of lineage specific markers, including those associated with transit amplifying cells, immature and mature enterocytes, and subsets of M cells, Paneth cells, goblet cells, and enteroendocrine cells (52–56), indicating a heterogeneous epithelial population. Low expression of markers such as *SI* suggests that our 2D iPSC-SIECs were less mature than gut-on-chip-cultured iPSC-SIECs (54,63), but 2D- iPSC-SIECs displayed broader epithelial lineage representation than previously reported Transwell-based iPSC-SIECs cultures (63). Overall, the data indicate that our protocol supports multilineage differentiation, while further optimization is needed to enhance enterocyte maturation and drug-metabolizing capacity.

Preceding studies have reported that the celiac intestinal epithelium contains a higher proportion of secretory cells, which are immature compared to absorptive enterocytes (77), both during active disease and after a prolonged adherence to a gluten-free diet (78). To investigate whether this reflects an intrinsic difference in differentiation potential, we compared the enterocyte gene expression between CeD-derived line and healthy control-derive line. While the expression of the enterocyte-associated genes *APOA1* and *DPP4* (52,53,56) showed minor variation between the lines, expression of the enteroendocrine marker gene *TTR* (52,53,56) was similarly expressed in SIEC-stage of both CeD-derived line and healthy control-derived line. Further cell deconvolution assay with Pan-GI Cell Atlas evinced that the epithelial cell type composition was comparable in both CeD-derived line and healthy control-derived line. This overall concordance supports the reproducibility of the 2D- iPSC-SIEC differentiation and maturation approach.

In active celiac disease elevated levels of IFNγ have been reported in the intestinal epithelium (28,30), accompanied by an upregulation of IFNγ-regulated genes (55,79). Dotsenko et al. (17) demonstrated that approximately 50% of the epithelial “gluten-induced genes” were also responsive to IFNγ. In the present study, we aimed to validate the gene expression of our 2D iPSC-SIEC *in vitro* model using IFNγ stimulation.

IFNγ stimulation did not alter overall epithelial morphology. Consistent with previous reports, IFNγ induced enhanced cytotoxic transcriptional signature than TNFα (80), similarly both in the CeD- derived line and the healthy-control-derived line. There were minor differences in transcriptomes between CeD-derived line and the healthy-control-derived line following the cytokine stimulation. To assess whether the observed differences were attributable to variations in cell type composition (54) a cell deconvolution analysis was conducted using the Pan-GI Cell Atlas. This analysis showed that there were no substantial differences in epithelial, mesenchymal or immune cell type composition between the CeD- derived line and healthy control-derived lines after cytokine stimuli. There might have been small differences in cell types and their differentiation, which would have reflected to this result, however, bulk RNA sequencing precludes definitive cell type-specific conclusions. Previous studies have demonstrated that donor variation is an important source of iPSC line-to-line heterogeneity (81,82). Moreover, even independently derived clones from the same donor may exhibit differences in genetic, transcriptional, and differentiation-related properties, highlighting the complex sources of variability inherent to iPSC-based models (83–85). Therefore, future application of this iPSC-SIEC model for the assessment of disease-specific functional phenotypes would require larger cohorts to robustly distinguish disease-associated effects from biological variability.

IFNγ robustly induced canonical interferon-responsive genes, including *STAT1*, *IDO1*, *CXCL9*, *CXCL10*, and *CXCL11*, in all iPSC-SIEC lines, in agreement with studies using biopsy-derived organoids (17), iPSC-derived organoids (36), and gut-on-chip models (54). Limited CXCL induction seen in previous reports e.g. Moerkens et al (54), likely reflect variations in stimulation duration and cytokine exposure.

Although bulk RNA sequencing precluded cell-type specific resolution, several IFNγ-responsive genes overlapped with those reported in organ-on-chip model (54), including *STAT1, IRF1, TAP1, WARS, PSMB9, IFIT3*, and *HLA E*, supporting shared epithelial response programs. Comparison with transcriptomic data from intestinal biopsies (86,87), and celiac patient-derived organoids (17) further revealed substantial overlap in IFNγ induced genes, such as *STAT1, TAP1, TAP2*, and *WARS1,* whereas little overlap was observed with blood-derived expression profiles (87). Together, these findings support the relevance of iPSC-SIECs as a model for studying epithelial specific IFNγ responses relevant to enteropathies, such as celiac disease pathogenesis.

Gene ontology analysis showed enrichment of IFNγ−responsive and MHC class I antigen processing pathways, consistent with previous intestine-on-chip studies (54), but with a broader range of affected pathways in 2D iPSC-SIECs. This likely reflects the longer IFNγ exposure used in this study (48 h), as cytokine stimulation beyond 24 h has been shown to induce more extensive de novo transcription, whereas longer exposures do not further enhance responses (36,88).

TNFα, which is elevated in active celiac disease and central to Crohn’s disease pathogenesis (31,32,89), activated TNF signaling in all iPSC-SIEC lines, as indicated by upregulation of TNF-related genes. To the best of our knowledge, only a limited number of studies have utilized human patient- derived organoids to investigate the effects of TNFα in the small intestine (90,91). TNFα stimulation have previously shown destroy tight junctions in iPSC-derived intestinal organoids (91). The overall epithelial morphology remained largely unchanged after TNFα stimulation. Tight junctions were not directly evaluated; however, the decreased lateral localization of phalloidin-stained actin observed across the CeD-derived, healthy control-derived, and T2D-derived cell lines may be indicative of changes in epithelial barrier organization. TNFα stimulation increased expression of inflammatory and immune associated genes, including CXCLs, HLAs, *IL32*, *IFI27*, *STAT1*, *WARS1*, *IDO1*, and *TAP1* without inducing apoptotic pathways. This aligns with reports from intestinal organoid models showing TNFα-driven inflammatory responses increase nectoptosis without apoptosis (90), supporting the physiological relevance of the 2D iPSC-SIEC system.

## 5. Conclusion

We established a streamlined 2D differentiation protocol that generates polarized iPSC-derived small intestinal epithelial cells (SIECs) without an intermediate organoid phase.

The CeD-derived cell line and healthy control-derived cell line iPSC-SIECs showed similar differentiation capacity. The IFNγ stimulation robustly activated canonical interferon pathways across all derived cell lines and recapitulated transcriptional signatures observed in celiac intestinal tissue, supporting the physiological relevance of the model.

Overall, 2D iPSC-SIECs provide a tractable platform for studying epithelial-specific immune responses in enteropathies, such as celiac disease, while further optimization and larger cohorts will be required to enhance maturation and generalizability.

## 6. Limitations

This study was based on single iPSC lines from CeD, healthy control, T1D, and T2D donors, limiting conclusions about interindividual variability and population level reproducibility. iPSC SIECs were differentiated directly on Transwell inserts without epithelial cell purification; therefore, it remains unclear whether additional purification would have improved culture maturity or cellular homogeneity. Differentiation and maturation were conducted in static 2D-conditions, and maturation may differ under dynamic organ-on chip culture. Finally, bulk RNA sequencing was used, which does not provide information on cell-type composition, transcript localization, or the proportion of cells expressing specific genes

## Author Contribution

Conceptualization, KJ-U; methodology, PKu, MAn, MKo, HKa, KTa, FSi, ESo, JKe, KLe, PSa, MNy, KA-S and KJ-U; validation, PKu, MAn, MKo, HKa, KTa, FSi and KJ-U; formal analysis, PKu, MAn, , HKa, MKo, KTa, FSi, ESo,NKa and KJ-U; investigation, PKu, MAn, HKa, MKo, KTa, FSi, ESo, NKa, SKa, JKe, KLe and KJ-U; resources, PSa, MNy, KA-S, KKa, KLi and KJ-U; data curation, PKu, MAn, KTa, HKa and KJ-U; writing/original draft preparation, PKu, MAn, HKa,MKo, KLi and KJ-U.; writing/review and editing, PKu, MAn, MKo, HKa, KTa, FSi, ESo, NKa, Ska, JKe, KLi, PSa, MNy, KA-S, KKa, KLi and KJ-U; visualization, MKo, HKa, PKu, MAn, PKu, KTa, FSi, and KJ-U; supervision, JKe, PSa, MNy and KJ-U; project administration KJ-U; funding acquisition, KJ-U, KLi, KKa, KA-S, MNy, PSa,. All authors have read and approved the published version of the manuscript.

## Institutional Review Board Statement

The derivation and expansion of iPSC lines was approved by the ethical committee of Pirkanmaa Hospital District (R12123), and written consent was obtained from all fibroblast donors. All patients were over 18 years of age. The usage of cell lines for gastrointestinal research was approved by the ethical committee of Pirkanmaa Hospital District (R20008). No new cell lines were derived in this study.

## Data Availability Statement

Requests for access to the detailed study data can be submitted to the Ethics Committee of the Wellbeing Services County of Pirkanmaa by email at or by post to Pirhan Tutkimuspalvelut, Eettinen toimikunta, PL 2000, 33521 Tampere, Finland.

## Acknowledgments

Soili Peltoniemi, Nina Koivisto, Markus Haponen, and MSc Sanni Erämies are thanked for their excellent technical assistance. The authors acknowledge Biocenter Finland (BF), Tampere iPSC core, Tampere Imaging Facility (TIF) core, and Tampere Genomics Facility core for their services.

## Conflicts of Interest

The funders had no role in the design of the study, in the collection, analyses, or interpretation of the data, nor in the writing of the manuscript or the decision to publish the results.

## Supporting Information

**S1 Fig. Differentiation of iPSCs towards DE, PDE and SIEC.** Before differentiation, both iPSC lines expressed the as well as the *Nanog* gene and protein (n = 3, (A), and *POU5F1* / Oct3/4 gene and protein (n = 3, (B). At the DE stage, cultures expressed the *FOXA2* and *SOX17* gene and protein ((C ) n = 3; FOXA2 red, SOX17 green, DAPI blue, 100 μm). At the PDE stage, cultures expressed the *Cdx2* and *SOX17* gene and protein ((D) n = 3; Cdx2 red, SOX17 green, DAPI blue, scale bar 100 μm). Microscope images (A-D) were taken with an Evident-Olympus IX83 microscope 20x lens using a Hamamatsu ORCA-Fusion C14440-20UP camera and NEO LiveImaging software. Intestinal epithelial cell specific functionality assessed with PEPT1 dipeptide uptake assay. On left there is image captured above the cellular level and on the right, there is image taken from the level which is in the middle of nucleus (n=3; dipeptide D-Ala-Leu-Lys-AMCA green, DAPI blue, 20 μm). Confocal images were taken with a Zeiss LSM780 laser scanning microscope with a 40x lens and ZEN Blue 3.6 software.

S2 Fig. Differentially expressed genes in iPSC- vs. DE-stage and DE- vs. PDE-stage. Scatter plots of differentially expressed genes (log2) in iPSC vs. DE-stage between the CeD-derived cell line and the healthy-control-derived cell line (A) and DE- vs PDE-stage between the CeD-derived cell line and the healthy-control-derived cell line (B).

**S3 Fig. Differentiation of iPSCs towards DE, PDE and SIEC.** The mRNAs collected at iPSC, DE, PDE and SIEC stage were analyzed with RNA sequencing, and results were separately plotted to heatmaps of the 50 most significantly altered genes (A) T1D-derived line and T2D-derived line, and to Venn diagrams (B).

**S4 Fig. Cellular morphology and gene expression, and cytotoxicity following cytokine stimuli**. Type 1 diabetic-derived line (T1D) and type 2 diabetic-derived line (T2D) iPSC-SIECs matured for 26 days were stimulated with IFNγ (10 ng/mL and 50 ng/mL) or TNFα (10 ng/mL) for 48 h. (A) Immunofluorescence staining for PEPT1 (green), filamentous actin (phalloidin, red), and nuclei (DAPI, blue). (B) SLC15A1 expression quantified by qRT-PCR. (C) Cytotoxicity is assessed with lactate dehydrogenase (LDH) activity in culture media after 48 h cytokine exposure. Statistical significance * = p ≤ 0.05; * = p ≤ 0.005, and *** = p ≤ 0.0005.

**S5 Fig. Differentially expressed genes after IFN**γ **stimulations.** Differentially expressed genes (log2) after 10 ng/mL and 50 ng/ml IFNγ stimulation, (A) in CeD-derived line, (B) in healthy control-derived line, (C) n type 1 diabetic-derived line, and (D) in type 1 diabetic-derived line.

**S6 Fig. Differentially expressed genes after TNF**α **stimulations.** Differentially expressed genes (log2) fter 10 ng/mL TNFα stimulation, (A) in CeD-derived line versus healthy control-derived line, (B) in T1D- derived line versus healthy control-derived, (C) in T2D-derived line versus healthy control-derived line.

**S7 Fig. Canonical_markers_heatmap depicting proportion of immune cells across** CeD-derived cell ine, healthy control-derived cell line, T1D-derived cell line and T2D-derived cell line in unstimulated, IFNγ 10 ng/mL and 50 ng/mL) or TNFα (10 ng/mL) stimulated iPSC-SIECs.

**S1 Table.** poly(A) capturing indexing oligos used for RNA sequencing library preparation

## Notes

### Competing Interest Statement

The authors have declared no competing interest.

### Summary of Updates

This manuscript is now entirely rewritten to temper the tone in celiac patient vs control patient comparisons. There is new data about transepithelial resistance measurements and enterocyte polarity (new figure 2), the figure 3 is now edited to be more clear, and new figure 9 to show different cell types in stimulated and IFNg stimulated samples. We have new supplemental figure to elucidate differentially expressed genes in iPSC-vs. DE-stage and DE-vs. PDE-stage, edited supplemental figure 3 to be more clear, added new supplemental figure 5 and 6 about differentially expressed genes after IFGg and TNFa ( respectively), and added new canonical markers heatmap showing the proportion of immune cells i samples, supplemental figure 7.

