## Supplementary figures and images for "Planar, organoid-independent iPSC-derived small intestinal epithelium from celiac disease patient for studying cytokine-induced changes"

### Supplemental Figure 1

Kukkoaho et al S1 Fig

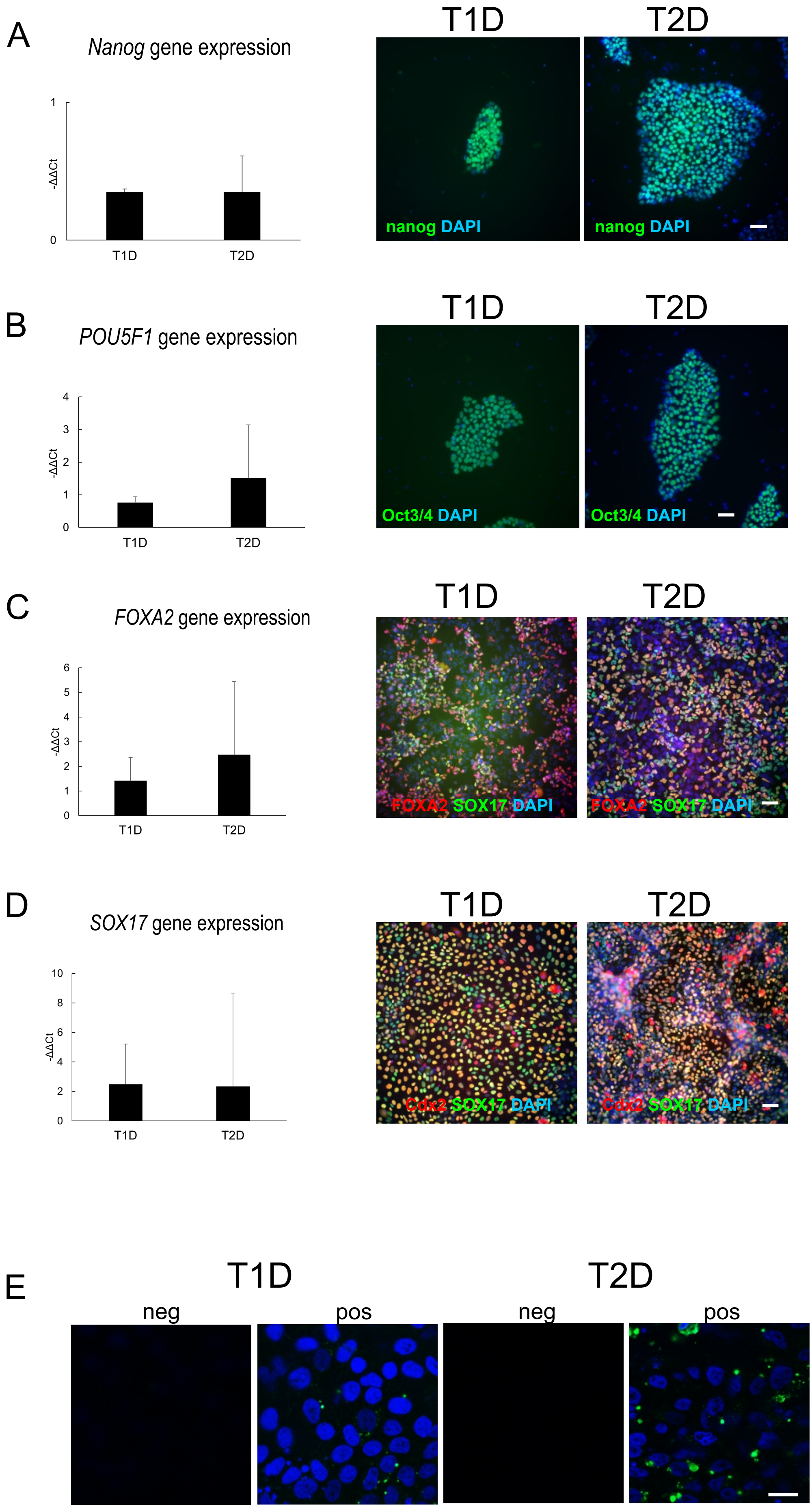

| p. adjust |
|---|
| 0,004 |
| 0,004 |
| 0,004 |
| 0,006 |
| 0,008 |
| 0,008 |
| 0,008 |
| 0,008 |
| 0,008 |
| 0,008 |
| 0,008 |
| 0,008 |
| 0,008 |
| 0,008 |
| 0,008 |
| 0,009 |
| 0,009 |
| 0,013 |
| 0,013 |
| 0,015 |

### Supplemental Figure 2

Kukkoaho et al S2 Fig

A

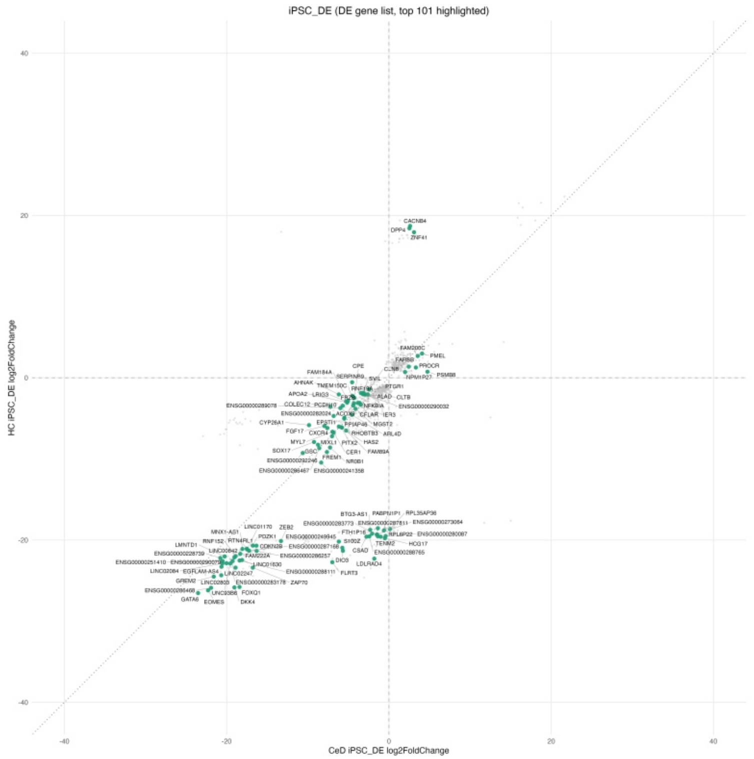


B

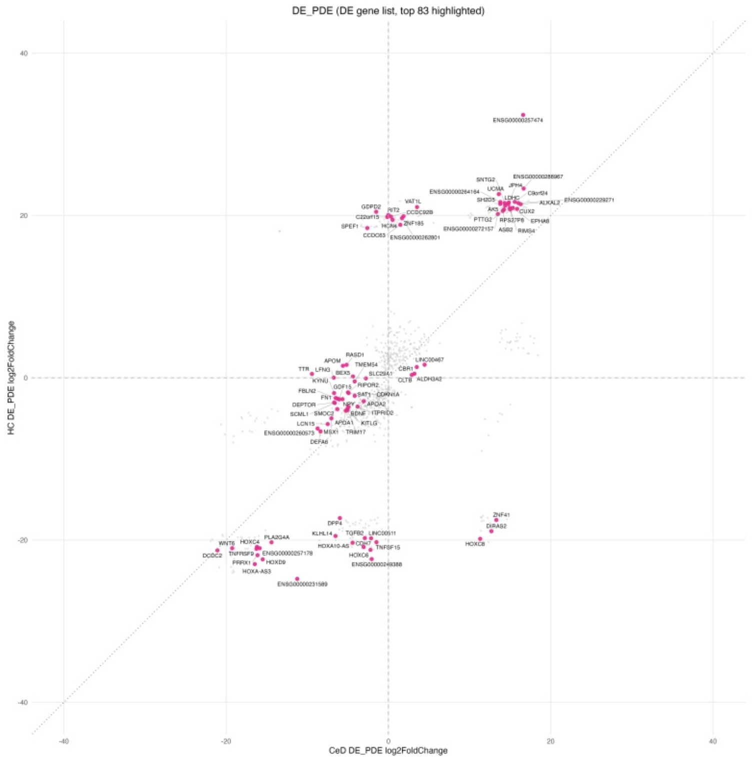

### Supplemental Figure 3

#### Kukkoaho et al S3 Fig

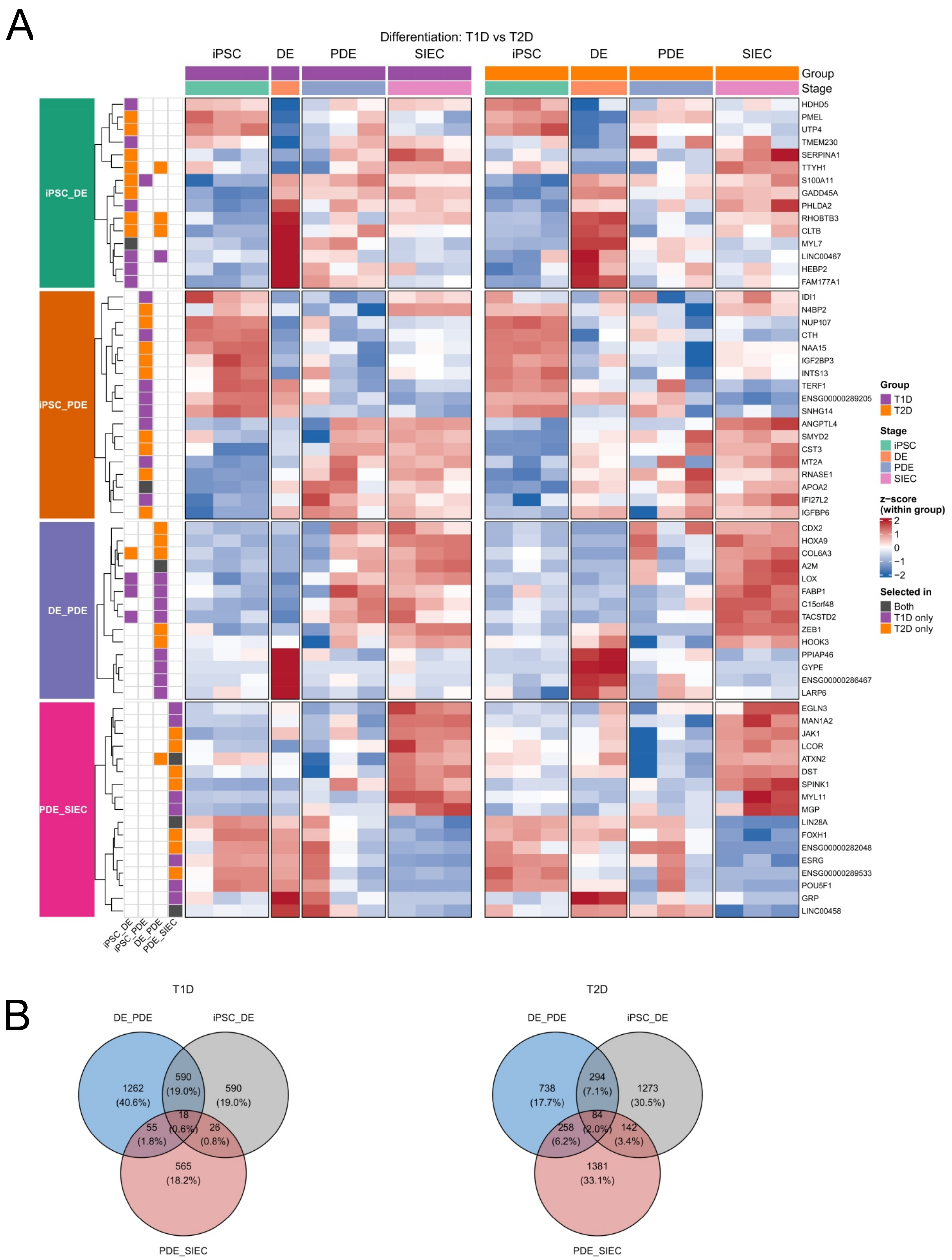

| p. adjust |
| --- |
| 0,004 |
| 0,004 |
| 0,004 |
| 0,006 |
| 0,008 |
| 0,008 |
| 0,008 |
| 0,008 |
| 0,008 |
| 0,008 |
| 0,008 |
| 0,008 |
| 0,008 |
| 0,008 |
| 0,008 |
| 0,009 |
| 0,009 |
| 0,013 |
| 0,013 |
| 0,015 |

### Supplemental Figure 5

Kukkoaho et al S5 Fig

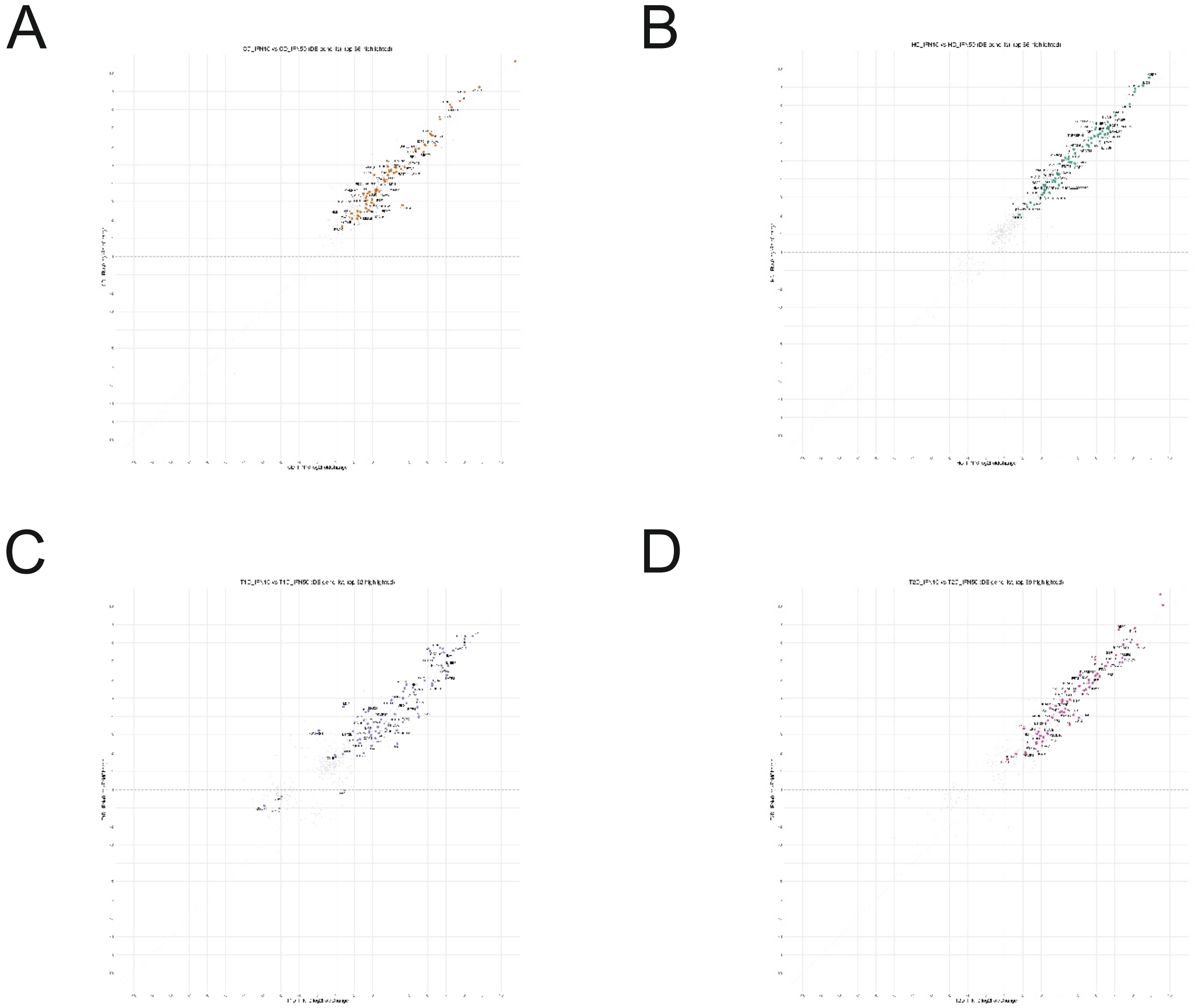

### Supplemental Figure 6

Kukkoaho et al S6 Fig

A

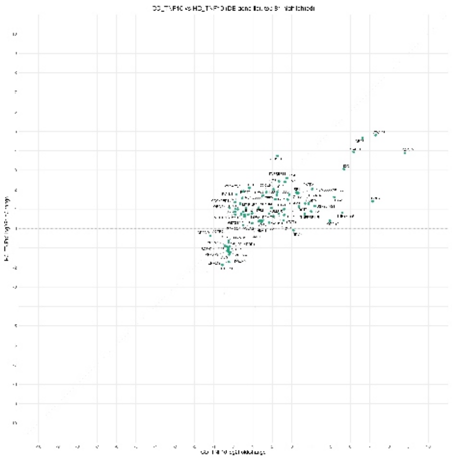

B

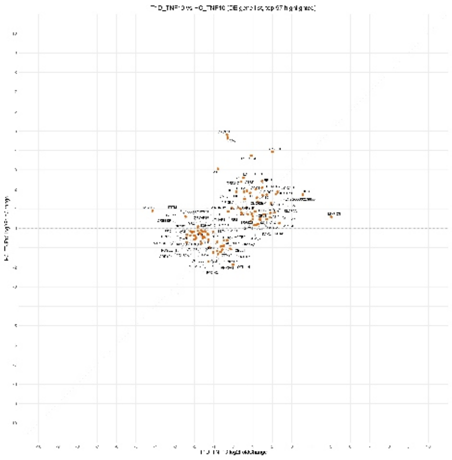

C

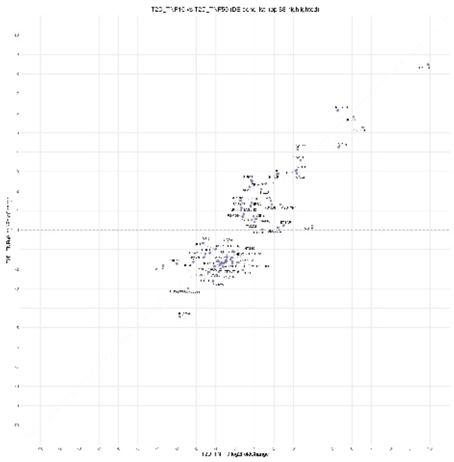

### Supplemental Figure 7

#### Kukkoaho et al S7 Fig

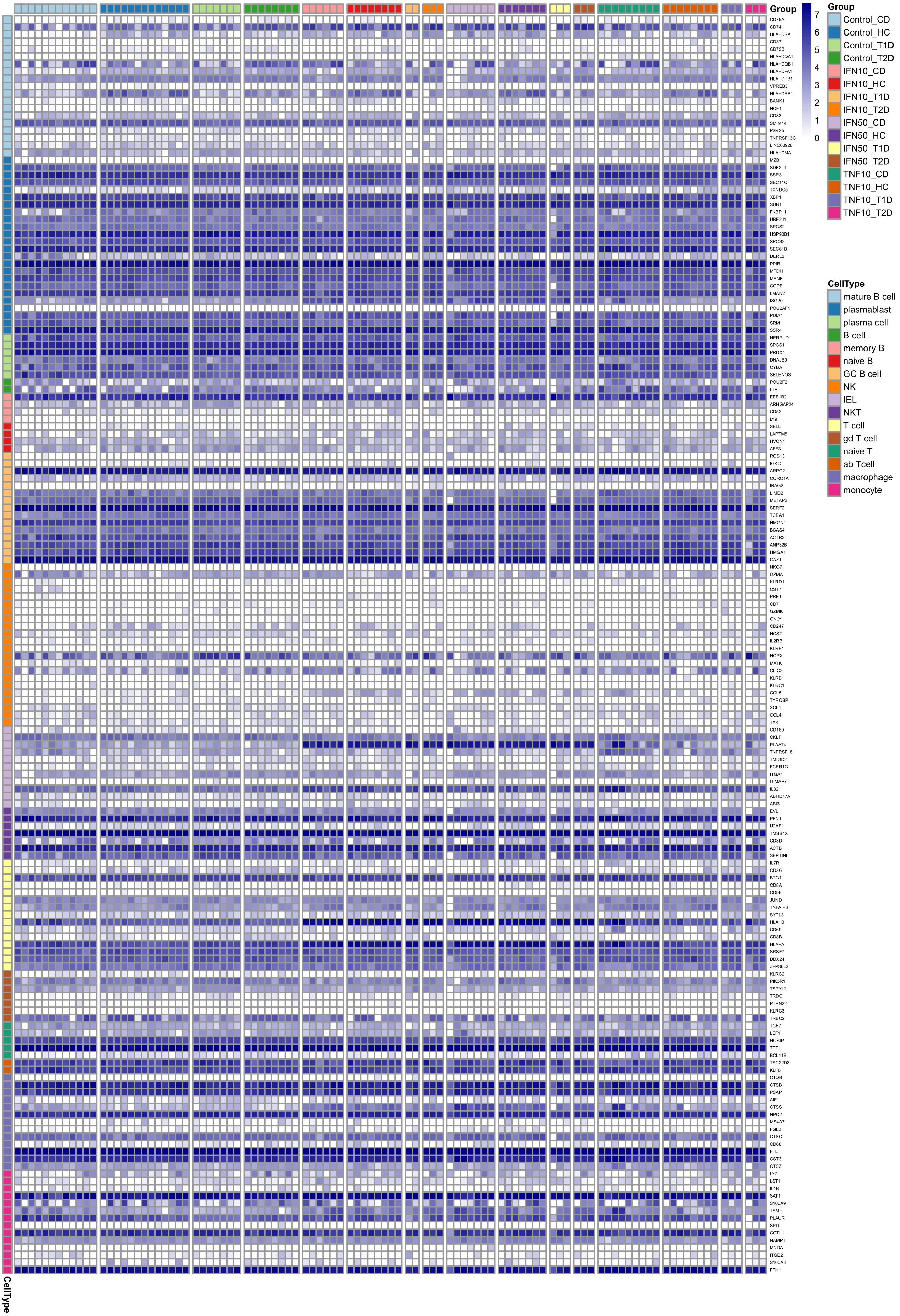
