## Supplemental Figure 4 for "Planar, organoid-independent iPSC-derived small intestinal epithelium from celiac disease patient for studying cytokine-induced changes"

A

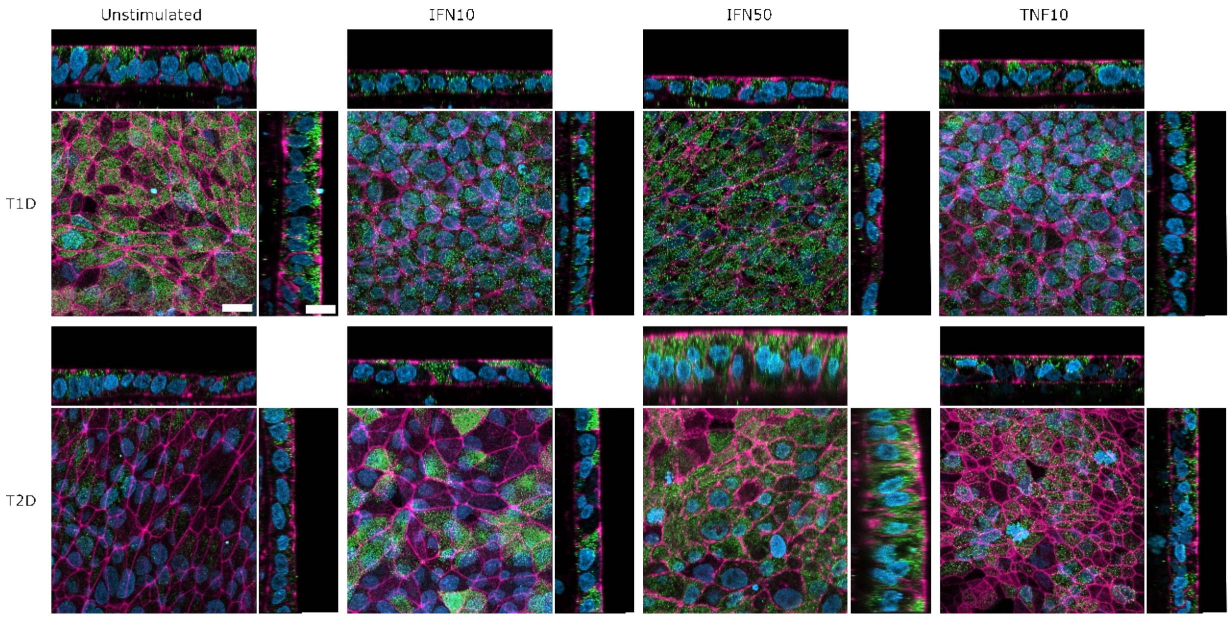
Unstimulated
IFN10
IFN50
TNF10
T1D
T2D

B

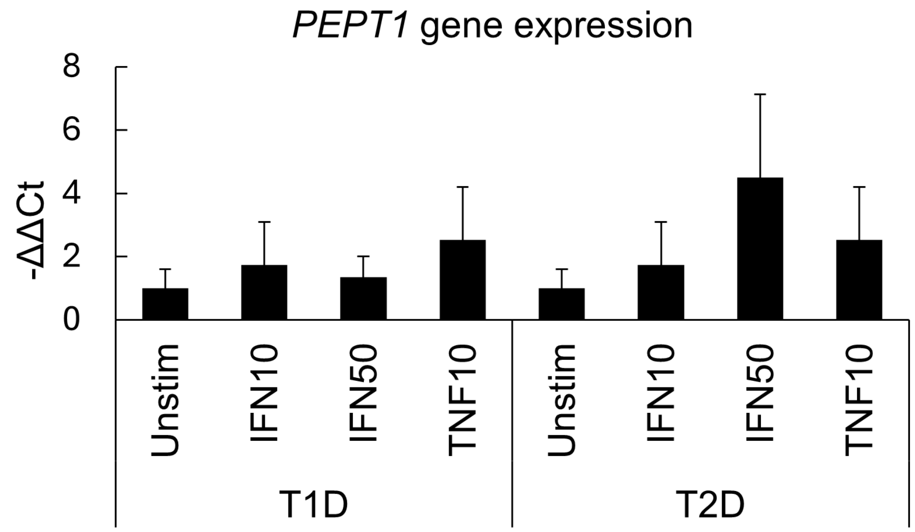
PEPT1 gene expression
-ΔΔCt
8
6
4
2
0
Unstim
IFN10
IFN50
TNF10
Unstim
IFN10
IFN50
TNF10
T1D
T2D

C

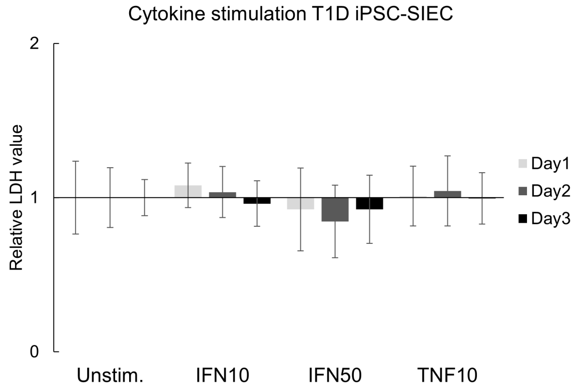
Cytokine stimulation T1D iPSC-SIEC
Relative LDH value
2
1
0
Unstim.
IFN10
IFN50
TNF10
Day1
Day2
Day3

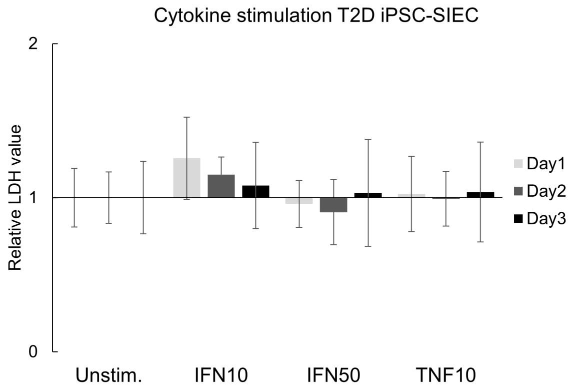
Cytokine stimulation T2D iPSC-SIEC
Relative LDH value
2
1
0
Unstim.
IFN10
IFN50
TNF10
Day1
Day2
Day3
